# Mechanical regulation of endocytosis by protein droplet capillary forces

**DOI:** 10.64898/2026.09.27.754800

**Authors:** Michael A. Ferrin, Dominic Mokbel, Marcel Mokbel, David G. Drubin, Sebastian Aland, Frank Jülicher, Tyler S. Harmon

## Abstract

Clathrin-mediated endocytosis (CME) is the primary pathway for internalization of extracellular and membrane cargo in eukaryotic cells. It is characterized by a patch of the plasma membrane invaginating and pinching off to generate a cytoplasmic vesicle. Recently, experimental evidence has begun to accumulate in support of proteins assembling into liquid-like condensates at nascent CME sites, but potential functional consequences are not understood. Given that condensate capillary forces can influence membrane bending, we constructed complementary analytical and numerical model systems to probe the mechanical influence of a protein droplet on endocytosis. Model evaluations and analysis reveal that a droplet can regulate the progression of endocytosis by stalling invagination until the droplet grows past a threshold size. After the droplet-mediated brake is released, the droplet provides a force that can drive the membrane bud toward scission. Thus, we propose a “mechanical timer” mechanism as a function for protein phase separation in cellular membrane trafficking.

**Statement of Significance:** Clathrin-mediated endocytosis is the essential pathway through which a patch of plasma membrane deforms into a vesicle in all eukaryotic cells. Recent experimental studies suggest that cytoplasmic components of the endocytic machinery condense into a liquid-like droplet on the membrane prior to internalization, raising questions about the droplet’s biological function. Using complementary numerical and analytical modeling approaches, we show that an endocytic droplet can regulate the pathway via capillary force. The droplet volume sets the strength of this force, providing the cell with a handle to switch from stalling to driving membrane invagination. While the biological role of liquid condensates is typically attributed to organizing biochemistry, this work demonstrates a novel purely mechanical function.

## Introduction

Capillary forces (also known as interfacial forces) are an intrinsic feature of systems undergoing liquid-liquid phase separation (LLPS).^1^ Surface tension drives the minimization of surface area of immiscible liquid phases, leading to a relaxation to circular shapes in 2D systems and to spherical droplet shapes in 3D systems. In the presence of a solid substrate or a third liquid phase, the relative magnitude of surface tensions among the three interfaces introduces an interfacial force balance that determines the droplet contact angle and governs wetting. As the catalog of coexisting membraneless compartments in living by cells formed by LLPS grows, it raises the exciting question whether the magnitude of their capillary forces is relevant to cellular physiology, and, if so, how they might be harnessed by biology.

Indeed, phase-separated droplet capillary forces are sufficient to drive a variety of membrane and droplet remodeling behaviors. These capillary forces have been experimentally observed across systems of varying complexity from minimal models composed of simple aqueous polymer droplets and phospholipid vesicles,^2–8^ to *in vitro* purified protein droplet and vesicle systems,^9,10^ even up to the living intracellular context.^11,12^

Membrane-associated protein droplets have been implicated in numerous intracellular membrane trafficking pathways, ^13–15^ including autophagy,^12,16^ presynaptic vesicle trafficking,^17–19^ endosome budding,^10,20^ endolysosome repair,^21^ secretory vesicle budding,^22^ Golgi processing,^23^ and endocytosis.^24–30^ The impact of capillary forces on the biology of these pathways remains largely unexplored.

Here, we investigate this question in the context of clathrin-mediated endocytosis (CME) in budding yeast. CME is an essential process for all eukaryotic cells in which a small patch of the cell membrane is pulled into a vesicle inside the cell. The pathway proceeds in a highly regular sequence of modular protein recruitment and membrane remodeling events (Figure 1 A). First, early-arriving proteins co-cluster with membrane cargo at nascent sites of endocytosis. They then recruit curvature-promoting proteins, which assemble into a clathrin lattice. The initiation phase has a variable duration, possibly due to a regulatory checkpoint.^28,31–34^ The assembly then transitions to a membrane-bending phase. Once the membrane gains curvature, forces generated by an active actin network pull the invagination inward. Finally, scission proteins are recruited and help pinch off an endocytic vesicle. Upon completion, the endocytic proteins recycle to the cytoplasm.

**Figure 1.**
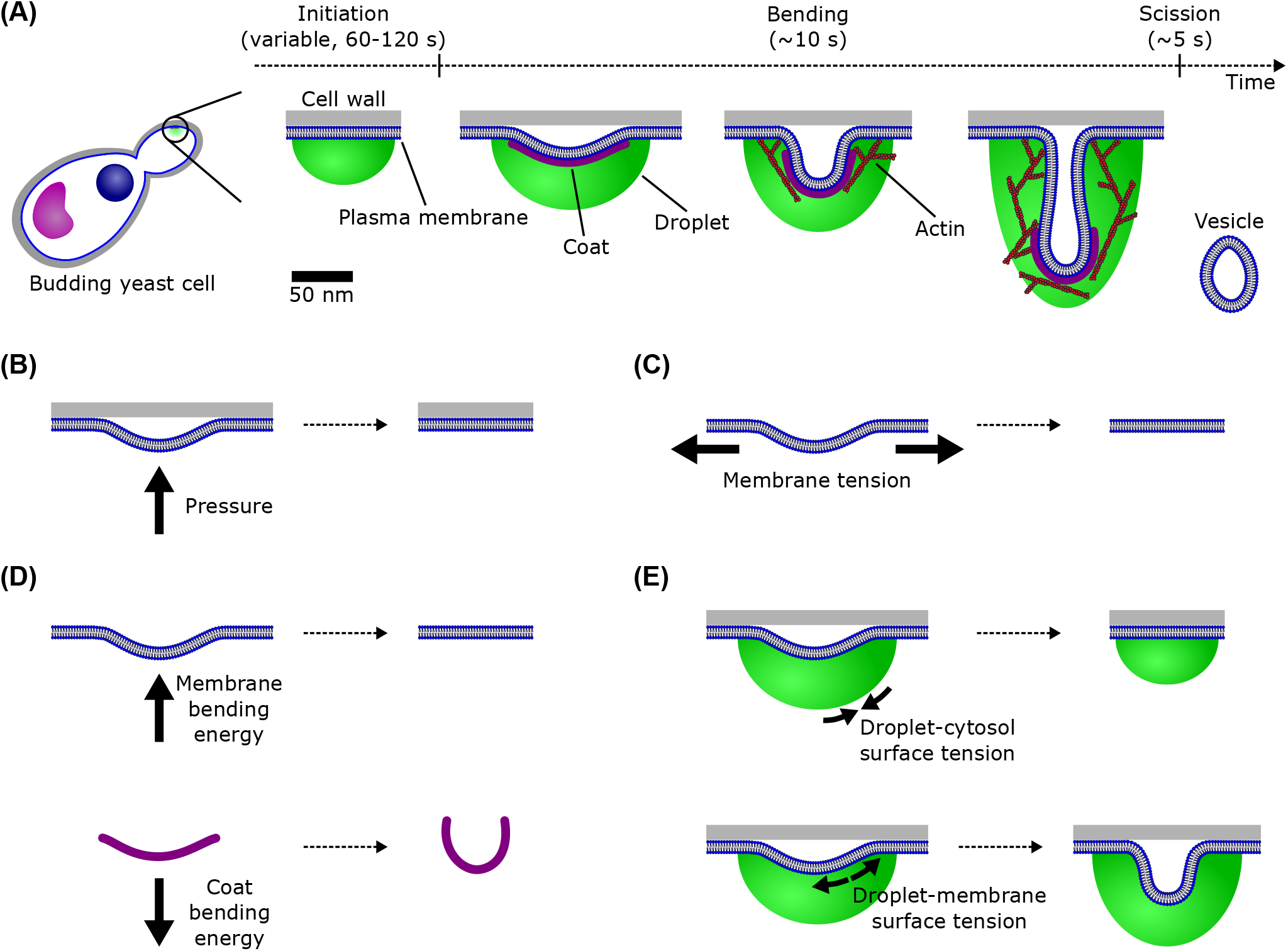
Endocytosis with a cytoplasmic protein droplet in budding yeast. (A) Clathrin-mediated endocytosis is a pathway for generating an intracellular vesicle by invaginating and pinching a patch of the plasma membrane while sequentially recruiting protein machinery from the cytoplasm. Following an early protein assembly phase with a highly variable lifetime among individual nascent endocytic sites, the pathway transitions to a membrane bending and scission phase with more regular dynamics. Early-arriving proteins are proposed to form a liquid-like droplet on the cytoplasmic face of the plasma membrane prior to membrane bending. (B-E) The mechanical forces impacting endocytosis with a droplet considered in this study are (B) turgor pressure forcing the membrane against the cell wall, (C) membrane tension resisting stretching the membrane, (D) bending energy from the membrane resisting increasing curvature and from the rigid protein coat driving increasing curvature, and (E) droplet surface tension driving it to minimize its contact area with the cytosol and increase its contact area with the membrane.

Budding yeast CME is excellently suited for studies of the biological impact of capillary forces. Several decades of detailed experimental investigations have yielded a comprehensive list of molecular participants and their dynamics. ^35,36^ More recently, evidence has arisen for early-arriving^24–26,37^ and late-arriving^37,38^ proteins forming droplets at endocytic sites, though the function of such droplets is under debate.

Using a combination of membrane wetting-coupled droplet dynamics numerical solutions and theory, we reveal a biological function for droplet capillary forces in endocytosis. We propose that a protein droplet can stall membrane bending during the initiation stage of endocytosis. This stalling force decreases as more droplet material is recruited, providing a natural timer to transition from the initiation to the membrane bending stage.

## Methods

We construct a simplified model of endocytosis represented in Figure 2 A, composed of a patch of heterogeneous membrane and an associated protein droplet. The droplet has a volume *V*_*d*_ and cytosol contact area *S*_*d*_. The membrane comprises three regions: the base surface contacting the cytosol *S*_*o*_, the base surface contacting the droplet *S*_*m*_, and the surface coated with curvature-generating proteins (such as clathrin) *S*_*c*_. The membrane invagination has the volume *V*_*c*_.

**Figure 2.**
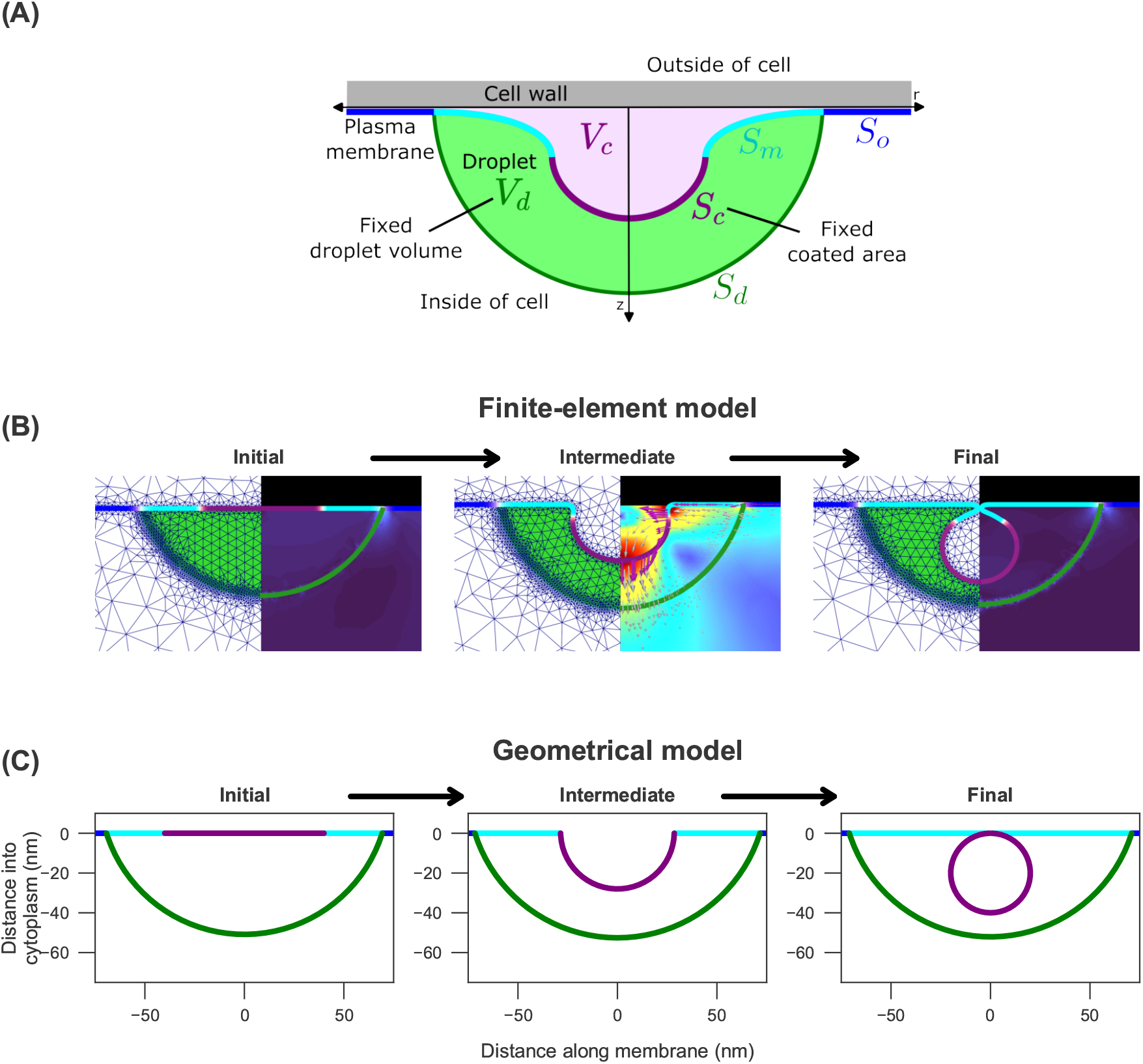
Schematic representation of modeling approaches. (A) Generic cartoon of yeast endocytosis highlighting the features shared between both model implementations. (B) Finite element model snapshots from three distinct time steps. The left side highlights the numerical grid with finer resolution at the droplet interface and at the membrane. The right side illustrates the velocity field, with the color code ranging from dark blue (slow) to red (fast). Black indicates zero-flow at the cell wall. (C) Cross-sectional snapshots of the geometrical model at different values of reaction coordinate h. P = 0.5 pN/nm^2^, σ = 0.2 pN/nm, γ_m_ = 3.2 pN/nm, γ_d_ = 10 pN/nm, 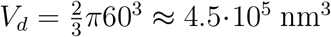, κ_c_ = 1500 pN·nm, C_c_ = 20^−1^ nm^−1^, S_c_ = 4π20^2^ ≈ 5 · 10^3^ nm^2^

The free energy *F* of the system is described by

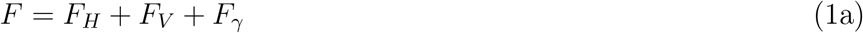

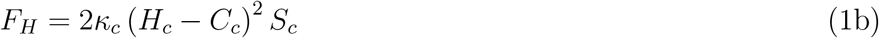

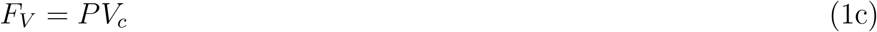

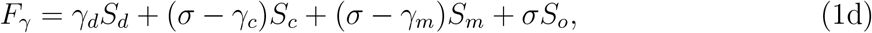

where *F*_*H*_ is the Helfrich bending energy of the membrane (Figure 1 D),^39^ *F*_*V*_ entails the turgor pressure opposing membrane invagination (Figure 1 B), and *F*_*γ*_ is the surface energy, composed of both membrane tension (Figure 1 C) and interfacial tensions from the droplet (Figure 1 E). Here, *κ*_*c*_ is the bending rigidity of the coated membrane, *H*_*c*_ is its mean curvature, *C*_*c*_ is its protein-induced spontaneous curvature, *P* is the fixed osmotic pressure across the membrane, and *σ* is the membrane tension which opposes stretching. Finally, the droplet contributes an interfacial tension *γ*_*d*_ associated with the droplet-cytosol interface *S*_*d*_, as well as adhesion energy densities *γ*_*m*_ and *γ*_*c*_ corresponding to the droplet-membrane interface *S*_*m*_ and the droplet-coat interface *S*_*c*_, respectively. We define both *γ*_*m*_ and *γ*_*c*_ as positive quantities; consequently, they enter the total surface energy with a negative sign, reflecting their adhesive character. The contact angle is determined by the balance of interfacial forces from Young’s equation 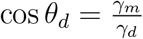.

The endocytic proteins experimentally demonstrated to form droplets (Ede1 in yeast^25^ and Eps15/Fcho1 in human^26^) arrive first in the stereotypical endocytic pathway, suggesting that they phase-separate before the curvature-generating coat is fully assembled. Consequently, by the time the coat is established and begins to drive membrane invagination, the droplet is already present and exerting capillary forces. To accurately capture this mechanical stage, our model initializes with both an established liquid droplet and a defined coated region on an initially flat membrane. Thus, the droplet volume *V*_*d*_ and coated membrane area *S*_*c*_ are considered as fixed quantities. This specific starting point allows us to investigate the direct mechanical competition at the onset of endocytosis: how the pre-existing capillary forces of the droplet interact with the coat-driven bending, and whether they are sufficient to mechanically resist or stall the invagination process.

We devise two distinct implementations of this model because each provides strengths that complement the limitations of the other. First, numerical solutions of a finite-element model provide accurate shapes of deformed fluid phospholipid membranes and protein droplets, as well as the evolution of the system over time. This numerical approach is complemented by a simplified geometrical model. This simplified model reveals interesting behaviors that can be explored over large parameter ranges. Solutions for critical points can be found analytically. This strength comes at the expense of morphological accuracy. Therefore, we combine both implementations to systematically study the key role of droplet capillary forces in membrane shaping proposed in this work.

### Finite element model

To determine the mechanical interplay between protein droplet wetting and membrane dynamics, we employ a continuum mechanics approach. Our finite-element computational framework builds upon the method introduced by Mokbel et al.,^54^ which combines a phase-field description of the bulk fluids (cytosol and droplet) with an Arbitrary Lagrangian-Eulerian (ALE) method for the explicit tracking of the deformable membrane shape *S*(*t*). The dynamic equations of the finite element model are obtained using the above free energy Equation 1, accounting for arbitrary deformation of the underlying surfaces. While the membrane is tracked as a two-dimensional curved surface, the droplet interface *S*_*d*_ is described by a phase field using an appropriately scaled Ginzburg-Landau energy functional. The resulting system couples Stokes hydrodynamics, capillary forces at all interfaces, and the bending mechanics of the membrane. Crucially, we extend this model^54^ to incorporate specific elements that play a key role in endocytosis: a spatially heterogeneous protein coat generating spontaneous curvature, an outward turgor pressure, and interactions with the rigid cell wall. Local area incompressibility of the membrane is enforced via a surface Lagrange multiplier field. To reduce computational cost while retaining 3D geometrical features, all computations are performed using an axisymmetric representation. Full details on the numerical implementation are provided in the Supporting Material.

### Geometrical model

To generate approximate solutions we study a simplified geometrical model. We constrain the geometry of the system to depend on a single degree of freedom: the normalized invagination distance *h*. The droplet shape is described as a spherical cap with a fixed volume and contact angle cos *θ*_*d*_ = *γ*_*m*_*/γ*_*d*_. Both base membrane areas *S*_*o*_ and *S*_*m*_ are constrained to be flat. The coated membrane region *S*_*c*_ is constrained to a spherical cap with fixed area that can transition from flat to spherical as the invagination distance *h* increases. The energy of a configuration is then given by Equation 1. As the coated membrane is a spherical cap, the mean curvature 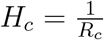, with *R*_*c*_ the spherical cap radius. These geometrical simplifications enable us to analytically compute the energy of conformations for given *h*.

## Results

### Droplet can stall membrane bending

Spatially heterogeneous spontaneous curvature and rigidity are known to drive a flat patch of membrane to deform into a nascent vesicle. ^55,56^ Our models reproduce this behavior in the absence of a droplet, which establishes a baseline for successful endocytosis (Figure 3 A, top, Movie S1). Such a morphological trajectory is consistent with previously published models of endocytosis.^40,57^ Upon adding a liquid droplet to the system, we observe a key change: instead of invaginating, the membrane deformation can stall early in the pathway (Figure 3 A, bottom, Movie S2).

**Figure 3.**
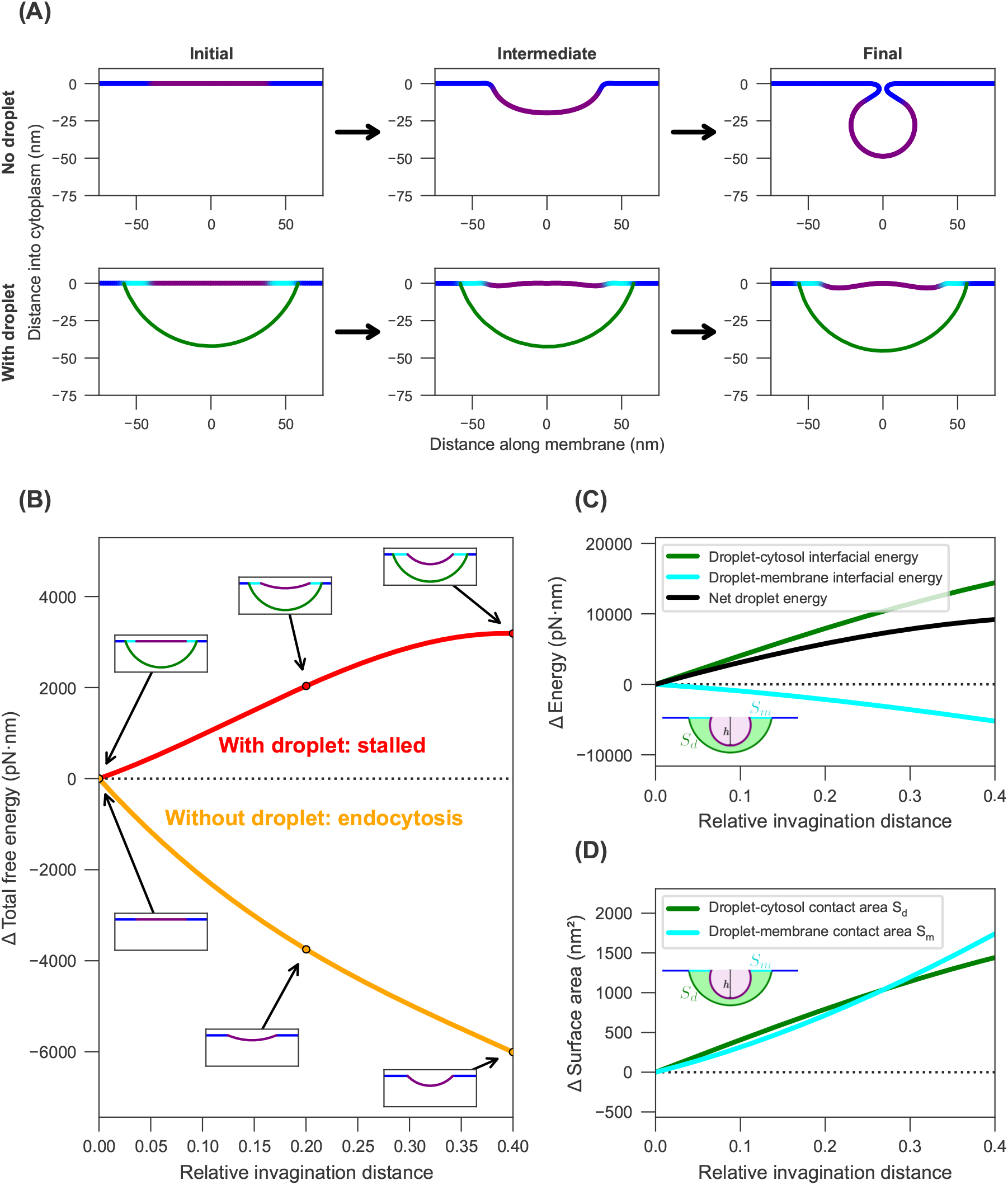
A droplet can stall the initiation of membrane bending. (A) Cross-sectional snapshots of endocytosis using the finite element model, showing the transition from the initial to the stationary state with no droplet (top) and with a droplet (bottom). (B) Plot of change in geometrical model total free energy Δ*F* over relative invagination distance *h* when droplet is present (red line) or absent (orange line). Cross-sectional snapshots for each model realization are shown as insets for *h* of 0, 0.2, and 0.4. Positive Δ*F* predicts stalling of membrane bending, while negative Δ*F* predicts spontaneous membrane bending. (C) Plot dissecting change in each geometrical model droplet energy term over *h*. Droplet-cytosol interfacial energy *γ*_*d*_*S*_*d*_ (green line) increases at a magnitude greater than droplet-membrane interfacial energy −*γ*_*m*_*S*_*m*_ (cyan line) decreases, resulting in a net increasing total droplet energy (black line); i.e., a droplet-mediated stalling force. (D) Plot of change in geometrical model droplet interfacial areas over *h*. Droplet-cytosol contact area *S*_*d*_ (green line) increases at a higher rate than droplet-membrane contact area *S*_*m*_ (cyan line) at *h* of 0, meaning that the droplet will always result in a net stalling force under the partial wetting condition *γ*_*d*_ *> γ*_*m*_. (A-D) *P* = 0.5 pN/nm^2^, *σ* = 0.2 pN/nm, *γ*_*m*_ = 3.2 pN/nm, *γ*_*d*_ = 10 pN/nm, *κ*_*c*_ = 1500 (A: 3173) pN · nm, *C*_*c*_ = 20^−1^ nm^−1^, *S*_*c*_ = 4*π*20^2^ ≈ 5 · 10^3^ nm^2^. (A bottom and B red) *V*_*d*_ ≈ 2.6 · 10^5^ nm^3^. (A top and B orange) *V*_*d*_ = 0 nm^3^.

We investigate the physical cause of this droplet-dependent stalling by exploring the geometrical model (see the Supporting Material and Figure S2). The geometrical model suggests that the flat coated membrane can spontaneously invaginate in the absence of a droplet, but a droplet wetting the surface can suppress the invagination (Figure 3 B). In this case, the free energy of the system with a droplet is at a local minimum at the flat initialization geometry.

Further analysis of the geometrical model reveals that a droplet will always introduce a resistive force to the start of endocytosis, independent of parameter choice. This is illustrated by isolating the droplet’s contribution to the free energy from Equation 1,

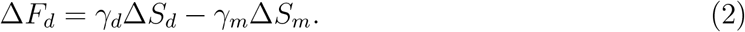

This equation indicates that the total change in energy from the droplet depends on the relative changes in droplet-cytosol contact area *S*_*d*_ and droplet-membrane contact area *S*_*m*_ (Figure 3 B-C). The *γ*_*c*_*S*_*c*_ term from Equation 1 drops out here due to our constant coated membrane area constraint, meaning that Δ*S*_*c*_ = 0. We represent the relative impact of these droplet energy terms in Figure 3 C. Despite wetting (*γ*_*m*_*S*_*m*_) providing an assistive force, the surface tension associated with the cytosolic interface (*γ*_*d*_*S*_*d*_) produces a net stalling force. This equation also highlights how the energy from the droplet directly relates to the change in each interface’s surface area, see Figure 3 D. The plot shows that *S*_*d*_ grows more quickly than *S*_*m*_ when membrane bending (*h* = 0) is initiated. More generally, the derivative of the droplet-cytosol and droplet-membrane contact area at the start of endocytosis are given by 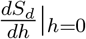 and 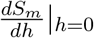, respectively. They behave relative to each other as

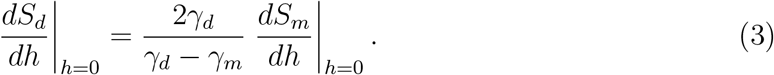

With the additional knowledge that *γ*_*d*_ *> γ*_*m*_ in order for partial wetting to occur (i.e., to give a droplet contact angle *θ*_*d*_ *>* 0 according to Young’s equation), 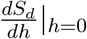 and 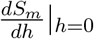 are always positive and 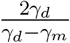 is always *>* 1.

This feature provides a general interpretation of the droplet mechanical energy from Equation 2: because *γ*_*d*_ *> γ*_*m*_ and Δ*S*_*d*_|_*h*=0_ *>* Δ*S*_*m*_|_*h*=0_, Δ*F*_*d*_|_*h*=0_ is positive. In other words, the droplet will contribute an increase in free energy at the beginning of endocytosis and will stall membrane bending.

### Droplet volume controls energetic barrier

Determining the change of *F*_*d*_ in Equation 2 for small *h*, to determine the droplet-mediated force at the initiation of endocytosis, gives

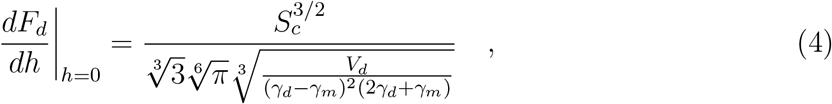

revealing that the resistive force from the droplet depends on the droplet volume *V*_*d*_ and its interfacial tensions *γ*_*d*_ and *γ*_*m*_. Interestingly, these are parameters that could vary over the course of endocytosis, by recruitment of droplet proteins and changing the overall protein composition of the droplet. The model predicts that the variation of one of these parameters alone can be sufficient to transition from a stalled to a permissive regime for membrane bending. We demonstrate this by plotting the solution of 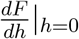 over the variation of these parameters (Figure 4 A). When the solution is positive (red shaded region), there is an energy barrier due to the droplet. When the solution is negative (green shaded region), the initial barrier is removed. We also show examples of free energy trajectories in a biologically relevant parameter range on either side of the transition point (Figure 4 B).

**Figure 4.**
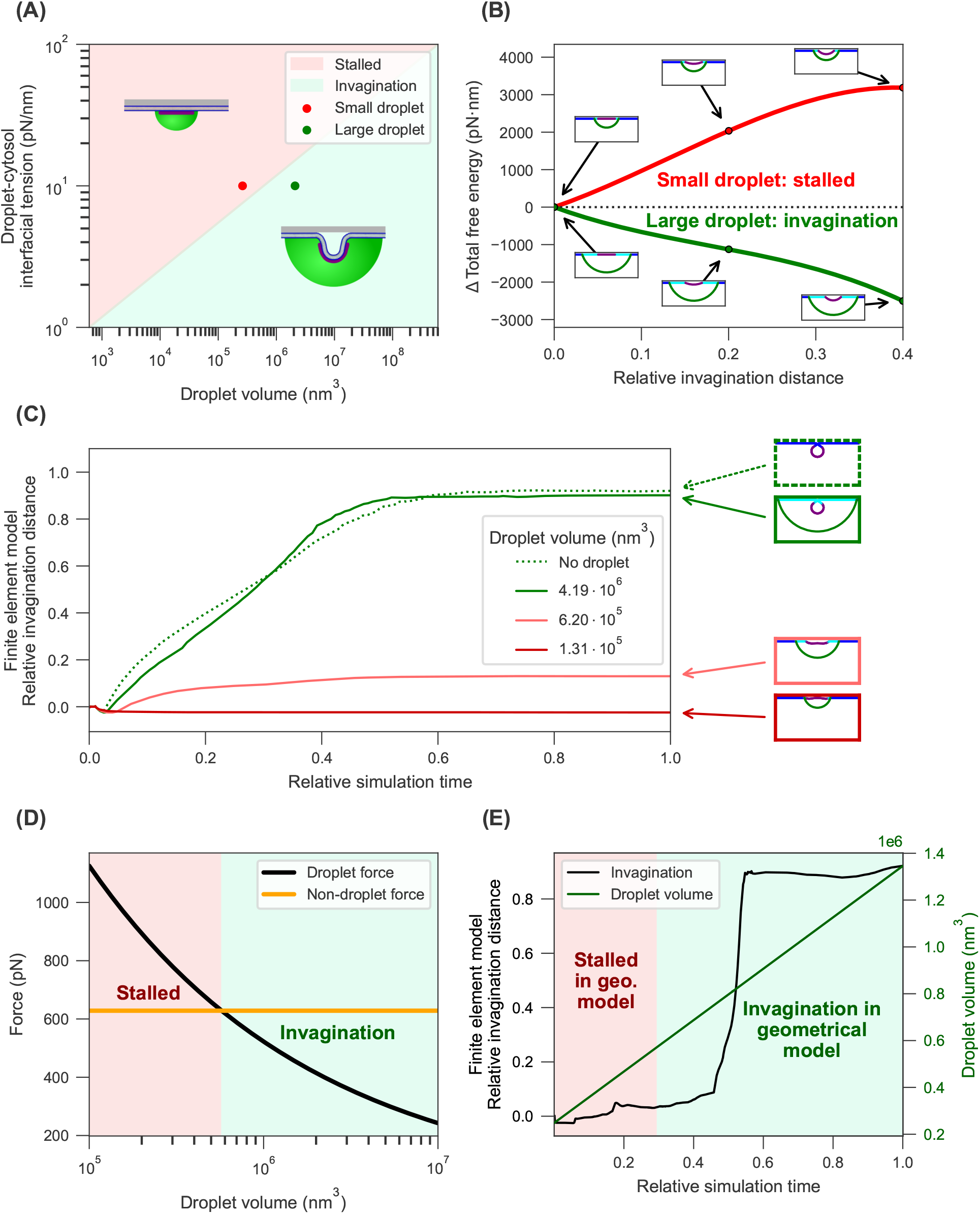
Droplet volume controls stalling force. (A) State diagram of geometrical model stalling behavior over droplet parameters. Stalling (red) denotes 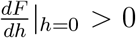; invagination (green) denotes 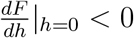. (B) Plot of change in geometrical model total free energy Δ*F* over invagination distance *h* at small droplet volume (red line, corresponding to red point in A) or large droplet volume (green line, corresponding to green point in A). Cross-sectional snapshots for each model realization are shown as insets for *h* of 0, 0.2, and 0.4. (C) Trajectories of numerical solutions initialized with different droplet volumes, or without a droplet. Snapshots of each solution at stationary state are shown to the right. (D) Plot of droplet volume (*V*_*d*_)-dependent stalling force (black line) compared to non-droplet force (orange line) in the geometrical model, both evaluated at the flat initiation state (*h* = 0). At small volumes (red shaded region), the droplet-mediated stalling force dominates. At volumes greater than a critical volume (green shaded region), the other forces dominate and invagination becomes spontaneous. (E) Numerical solution invagination trajectory (black line) where an initially small droplet increases in volume at a constant rate over time (green line). For comparison to the geometrical model, the time before reaching the predicted critical volume is the red shaded region of the plot, and the time after passing the critical volume is shaded green. (A,B,D) *P* = 0.5 pN/nm^2^, *σ* = 0.2 pN/nm, *κ*_*c*_ = 1500 pN · nm, *C*_*c*_ = 20^−1^ nm^−1^, *S*_*c*_ = 4*π*20^2^ ≈ 5 · 10^3^ nm^2^. (A) *γ*_*m*_ = 0.32*γ*_*d*_ in order to fix droplet contact angle *θ*_*d*_ (B,D) *γ*_*m*_ = 3.2 pN/nm, *γ*_*d*_ = 10 pN/nm (B, red) *V*_*d*_ ≈ 2.6 · 10^5^ nm^3^. (B, green) *V*_*d*_ ≈ 2.1 · 10^6^ nm^3^. (C, E) With the exception of *V*_*d*_, the parameters are given in Table 1.

To further test this control over stalling by droplet size in a physiological range, we computed numerical solutions that were initialized with different droplet volumes (Figure 4 C). As predicted by the geometrical model, small droplets stall invagination, while large droplets result in invagination dynamics similar to solutions with no droplet.

Finally, we can connect the droplet size-dependent stalling to cellular behavior via droplet growth dynamics. The geometrical model predicts a critical droplet volume at which the droplet stall force is overcome by the bending forces from the rest of the system (Figure 4 D, Movie S3). Numerical solutions with a time-dependent droplet volume also exhibit stalled membrane dynamics until passing a threshold volume that triggers rapid invagination (Figure 4 E, Movie S4). These numerical results are consistent with stereotypical observations of endocytic dynamics in live cells via time-lapse fluorescence microscopy. Putative droplet-forming proteins gradually accumulate at sites of endocytosis until they cross a threshold concentration, which triggers membrane internalization. ^58^

To place these forces in the context of the mechanical landscape at a yeast CME site, we report each of the individual force contributions in the geometrical model in physical units of pN (Figure S4). We compute forces as the derivative of the free energy with respect to the physical invagination distance *L*_*c*_ (Equation S23), such that the quantities shown are directly comparable to force values reported for other components of the CME machinery. For our reference parameter values, the droplet stalling force at the flat initiation state (*h* = 0) is of the order 10^2^ − 10^3^ pN (Figure S4 A), on the same scale as turgor pressure force resisting invagination and actin-derived assistive force (~ 800 - 2400 pN across budding and fission yeast estimates^59,60^). See Supporting Material section *Free energy and forces* for the derivations of these forces in the geometrical model.

### Droplet can drive scission

Although the droplet resists membrane bending at the initiation of endocytosis, its mechanical effect reverses as invagination progresses. At greater invagination distance, the geometrical model predicts that the droplet instead assists membrane deformation, driving the system toward completion of endocytosis (*h* = 1, corresponding to formation of the final endocytic vesicle via scission) (Figure 5 A-B). This switch from stalling to assisting occurs because both droplet interfacial energy terms decrease late in the pathway, resulting in a net assistive force from the droplet (Figure 5 B). Decomposing the droplet energy *F*_*d*_ to its relevant geometrical terms once again with Equation 2, we observe how this assistive droplet force arises from a decreasing droplet-cytosol contact area *S*_*d*_ and increasing droplet-membrane contact area *S*_*m*_ at high invagination distance (Figure 5 C).

**Figure 5.**
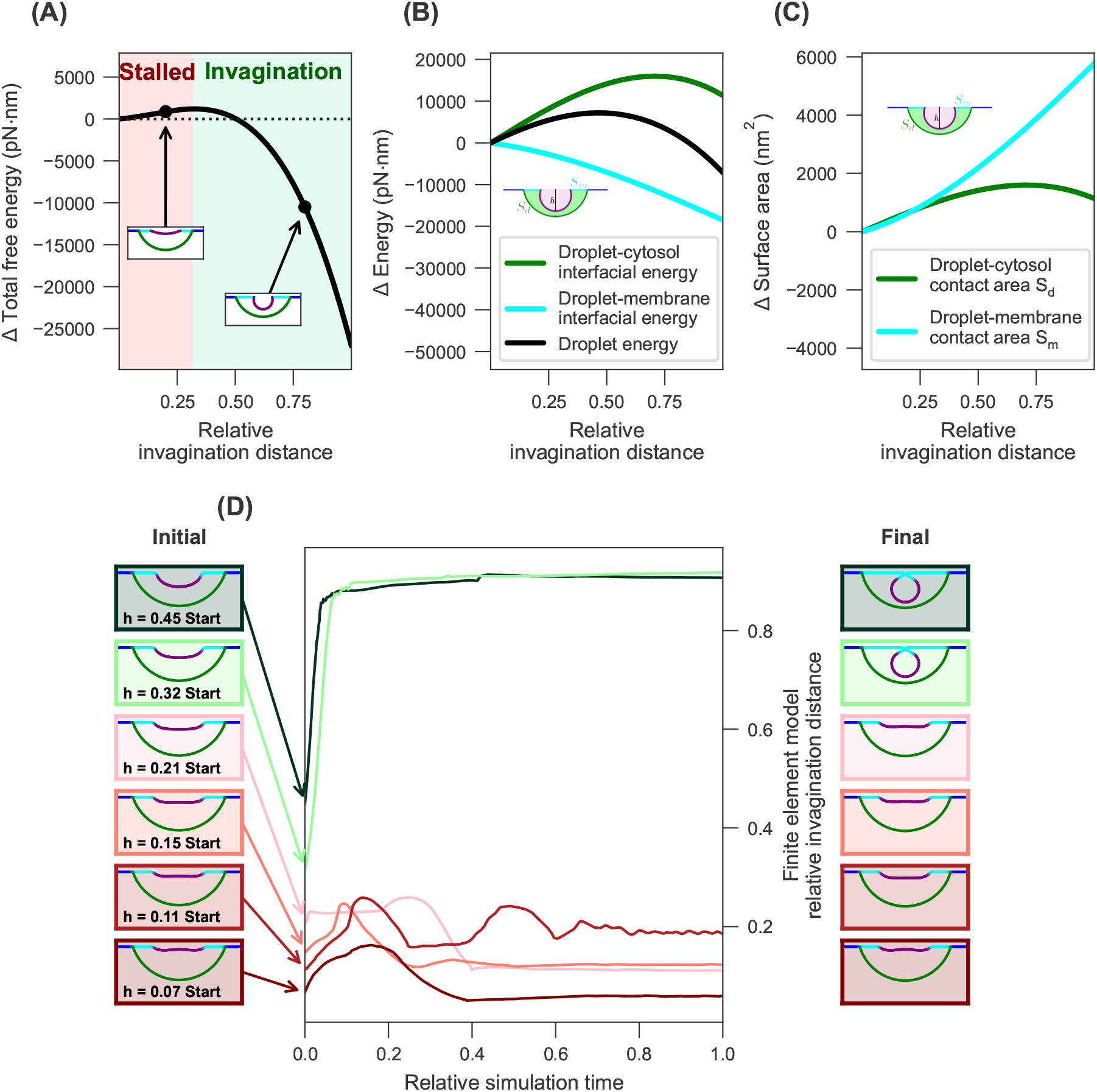
Droplet assists membrane invagination after energy barrier. (A) Change in geometrical model total free energy Δ*F* over full range of invagination distance *h*. Invagination is stalled up to the energy barrier at *h* of 0.39 (red shaded region), after which invagination is kinetically spontaneous (green shaded region). Cross-sectional snapshots are shown as insets for *h* of 0.2 and 0.8. (B) Change in each droplet energy term over *h*. Droplet-cytosol interfacial energy *γ*_*d*_*S*_*d*_ (green line) reaches a maximum at high *h*, while droplet-membrane interfacial energy −*γ*_*m*_*S*_*m*_ (cyan line) only decreases. This results in a barrier in the total droplet energy (black line); i.e., droplet-mediated stalling at low *h* and droplet-mediated invagination at high *h*. (C) Change in droplet interfacial areas over *h*. Droplet-cytosol contact area *S*_*d*_ (green line) reaches a maximum at high *h*, while droplet-membrane contact area *S*_*m*_ (cyan line) only increases, meaning that the droplet will always result in a net invagination force beyond a barrier *h*. (D) Trajectories of numerical solutions initialized at different intermediate invagination morphologies. Initial snapshots corresponding to each trajectory are shown to the left of the plot, and stationary state snapshots to the right. (A-C) *P* = 0.5 pN/nm^2^, *σ* = 0.2 pN/nm, *γ*_*m*_ = 3.2 pN/nm, *γ*_*d*_ = 10 pN/nm, *V*_*d*_ ≈ 2.6 · 10^5^ nm^3^, *κ*_*c*_ = 1500 pN nm, *C*_*c*_ = 20^−1^ nm^−1^, *S*_*c*_ = 4*π*20^2^ ≈ 5 · 10^3^ nm^2^ (D) Values of *V*_*d*_ for the snapshots, from bottom to top ≈ (4.77, 4.73, 4.68, 4.61, 4.5, 4.4) · 10^5^ nm^3^. The remaining parameters are listed in Table 1.

The geometrical model reveals how general the droplet-driven scission effect is. The change in droplet-cytosol surface area *S*_*d*_ at the end of membrane invagination is given by

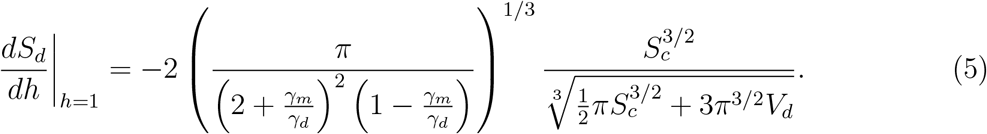

Again considering that partial wetting dictates *γ*_*m*_ *< γ*_*d*_, this complete expression always evaluates to a negative number. Thus, the energy from the droplet-cytosol interface will always decrease at late invagination because the interfacial area is decreasing.

The accompanying change in droplet-membrane surface area *S*_*m*_ is given by

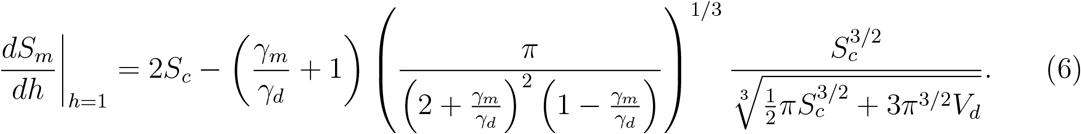

The sign of this expression depends on parameter choice, rather than conveniently evaluating as always positive or negative. However within a biologically relevant parameter range, this does result in a positive area change (Figure S3), meaning that this droplet interface also contributes an assistive force at late invagination.

We complemented this geometrical model analysis with numerical solutions initialized at different intermediate morphologies along the non-droplet internalization trajectory (Figure 5 D). The system is initialized by applying a uniform, standardized droplet phase-field profile across all membrane geometries. This baseline profile corresponds to a nominal droplet volume of approximately 5 · 10^5^ nm^3^ on a flat membrane. Because the membrane invagination depth varies across the setups, superimposing this fixed initial profile results in slightly different effective enclosed volumes, with a maximum deviation of up to 8% (see Figure 5 D). Crucially, the respective effective droplet volume is strictly conserved during the dynamic evolution of each individual solution, ensuring that the mechanical droplet-membrane interplay is accurately captured. Here we also identify a droplet-induced energy barrier: the membrane stalls if initialized at a relative internalization distance of less than 0.32. Yet if initialized at an intermediate geometry beyond this stall boundary, the system rapidly goes to the final vesicle scission state.

## Discussion

Taken together, our theoretical approach suggests a mechanical function for phase-separated protein droplets in endocytosis (Figure 6). We propose that the first-arriving proteins in the pathway condense into a small droplet on the plasma membrane, consistent with recent purified protein and live cell imaging studies.^25,26^ The formation of the droplet interface immediately generates an opposing force to endocytic membrane bending, allowing cargo and curvature-generating proteins to accumulate without prematurely triggering internalization. The droplet-mediated stalling force becomes weaker as more proteins are recruited from the cytoplasm, and eventually gets overtaken by the coat’s curvature-generating force. After the nascent vesicle begins budding, the droplet switches from resisting to further driving the membrane internalization. Thus, the droplet serves to mechanically regulate endocytosis and provide a timer for the initiation and completion of endocytosis.

**Figure 6.**
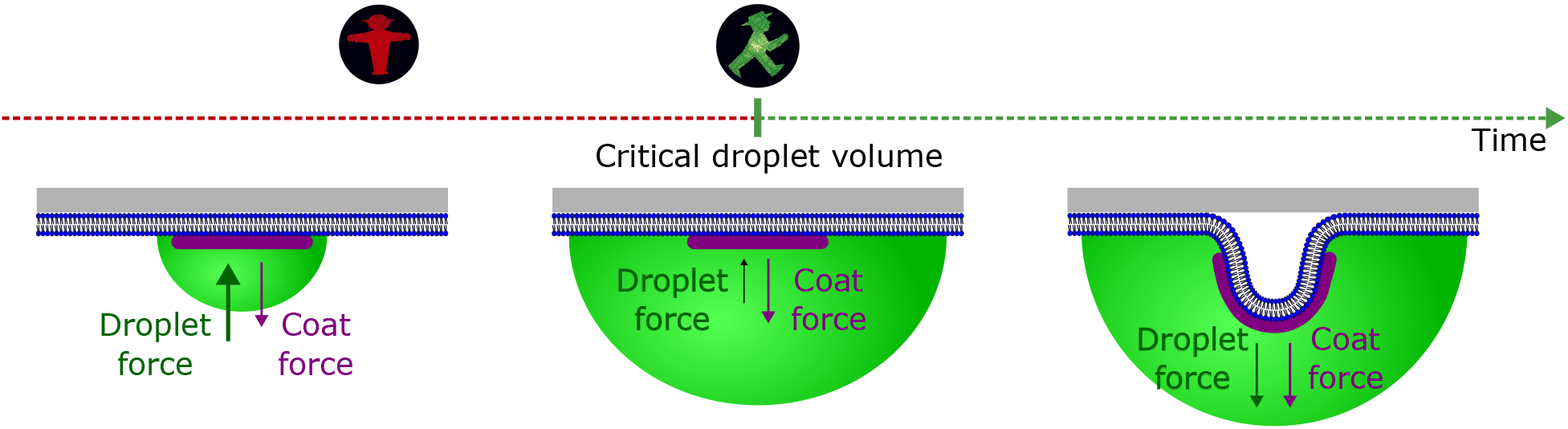
Conceptual model of droplet-mediated regulation of endocytosis. Our theory suggests a regulatory mechanism in which condensation of early endocytic proteins into a membrane-associated droplet produces a force against the membrane that stalls membrane bending. As cargo clustering drives recruitment of more proteins from the cytoplasm, the droplet increases in volume until it reaches a critical threshold where the stalling force is overcome by curvature-generating coat forces. After membrane bending is initiated, both the coat and droplet robustly drive invagination to completion.

In order to study general principles of how droplets impact the endocytic pathway, we approximated the endocytic protein network by an average composition. This considers the properties of the membrane, coat, and droplet at a bulk level. It is known that the recruitment of different proteins occurs at different times which would change the droplet composition over time.^37^ This feature has the potential to impart temporal control over the surface tension and wetting energies. We explicitly showed that droplet growth could relieve endocytosis stalling, but our results show that stalling could instead be relieved by changing surface interaction strengths (Figure 4 A).

In this work we focused on forces derived from thermodynamic interactions between components and surfaces and how these impact endocytosis. However, it is known that active forces from actin and myosin are essential for yeast clathrin-mediated endocytosis under normal laboratory conditions.^61,62^ We compensated for the lack of active forces in our models by increasing the assistive force from the clathrin coat via a higher spontaneous curvature than previous models of yeast endocytosis.^48,50^ The absence of axial pulling forces as well as the high preferred curvature could explain why our computed trajectories progress through morphologies resembling a stereotypically mammalian spherical intermediate ^63,64^ rather than a tubular intermediate that is commonly observed in budding yeast. ^49^ However, we demonstrate here that the droplet performs its most important mechanical function during the flat-membrane initiation stage, before the actomyosin network arrives in the typical pathway.^65^ Our main conclusions regarding the droplet’s role as an energetic barrier therefore remain valid under these potentially different late-pathway forces and geometries.

Our model design was primarily inspired by yeast clathrin-mediated endocytosis, which raises the question whether it can be generalized to other species. The yeast cell wall provides a strong opposing force to bending the plasma membrane towards the outside of the cell. In contrast, a mammalian plasma membrane without a cell wall may bulge outward as a result of droplet capillary forces. It will be interesting to investigate in the future whether other structures of mammalian cells such as the actin cortex could also provide similar stabilizing forces, and what range of droplet interfacial tension could result in a realistic early endocytic membrane profile.

Finally, the impact of droplet interfacial tension on endocytic stalling (Figure 4 A) suggests that it is an important parameter to estimate and to perturb experimentally. Interfacial tension measurements have been reported for only a small catalog of biomolecular condensates,^45^ limiting our ability to constrain this model parameter. The value of the baseline droplet-cytosol interfacial tension (*γ*_*d*_) of 10 pN/nm lies within an order of magnitude of previously measured surface tension in polypeptide-nucleic acid condensates *in vitro*.^44^

More broadly, our work could help bring the fields of membrane trafficking cell biology and condensate biophysics closer together, and offer a new perspective at their intersection. A long-standing mystery in the mechanisms of endocytosis is the molecular basis for a proposed “checkpoint” that regulates the transition from the early protein accumulation stage to the late invagination stage.^31,32,34^ The droplet-mediated stalling mechanism presented here could play such a role. So far, suggested functions of condensates in cells have been typically the regulation of biochemical reactions and the positioning of biochemistry in the cell. The mechanical regulation of endocytosis proposed in this work expands possible functions of condensates and extends the field studying condensate capillary forces.^1^ With mechanical force generation added to the list of suggested functions of condensates, we are eager to explore how other membrane remodeling pathways may harness this potential.

## Author contributions

M.A.F., D.G.D., F.J., and T.S.H. conceived the project. M.A.F. and T.S.H. developed and analyzed the geometrical model. D.M., M.M., and S.A. developed the numerical model. D.M. implemented the numerical solver, performed simulation studies, and analyzed numerical data. M.A.F., D.M., D.G.D., S.A., F.J., and T.S.H. discussed the results and wrote the manuscript. D.G.D., F.J., and T.S.H. funded and supervised M.A.F. S.A. funded and supervised D.M. All authors approved the final version.

## Acknowledgement

This work was supported by the NSF Graduate Research Fellowship Program to M.A.F. T.S.H. acknowledges support from the German Research Foundation (DFG) under “Biomolecular Condensates: From Physics to Biological Functions” (RTG 3120). S.A. acknowledges support from the DFG (grants 422878782 and 570692780). This research was supported in part by grant NSF PHY-2309135 and the Gordon and Betty Moore Foundation Grant No. 2919.02 to the Kavli Institute for Theoretical Physics (KITP). D.G.D. was supported by NIH MIRA grant R35GM118149. The authors gratefully acknowledge computing time on the high-performance computer at the NHR Center at TU Dresden. This center is jointly supported by the Federal Ministry of Education and Research and the state governments participating in the NHR.

## Supporting material

### Details on the numerical method

To investigate the mechanical regulation of endocytosis, we employ a continuum mechanics approach that couples the fluid dynamics of the bulk phases with the mechanics of the lipid membrane. The system consists of two bulk phases (cytosol and protein droplet) separated by a diffuse interface, and a sharp interface representing the membrane *S*(*t*).

The methodology described below is based on the framework developed by Mokbel et al.^54^

### Fluid dynamics and phase separation

The bulk domain contains two immiscible fluids described by a phase-field variable *ϕ*, varying continuously from *ϕ* = 0 (cytosol) to *ϕ* = 1 (droplet) across an interface of width *ϵ*. The driving force for phase separation is the minimization of the bulk free energy 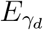, given by the Cahn-Hilliard functional:

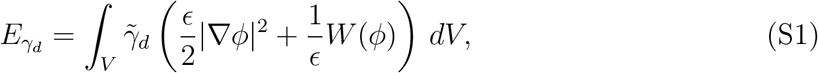

where 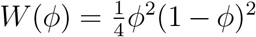 is a double-well potential and 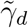 scales with the physical surface tension 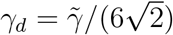.

The hydrodynamic evolution is governed by the incompressible Navier-Stokes equations coupled to the Cahn-Hilliard kinetics. The fluid velocity **u**, pressure *p*, and chemical potential *µ* evolve according to:

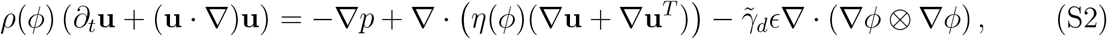

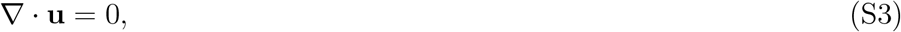

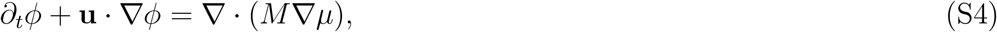

where 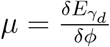 denotes the chemical potential and *M* denotes the mobility (see the section on weak formulations below for details). The density *ρ*(*ϕ*) and viscosity *η*(*ϕ*) interpolate linearly between the phase values. Note that in this work we assume Stokes flow and therefore set *ρ*(*ϕ*) = 0.

### Membrane mechanics and coupling

The membrane *S* moves with the local fluid velocity. Its mechanical response is governed by the Helfrich bending energy *E*_*H*_ and a tension energy 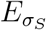. To model the biological reality of clathrin-mediated endocytosis, we introduce a spatially heterogeneous protein coat using a surface indicator function *ϕ*_*S*_ defined on *S* (*ϕ*_*S*_ ≈ 1 for coated, *ϕ*_*S*_ ≈ 0 for bare membrane). The total membrane energy is:

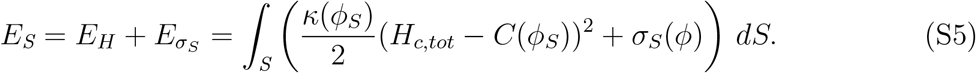

Here, *H*_*c,tot*_ is the sum of the principal curvatures (total curvature). Note that *E*_*H*_ is equal to the energy *F*_*H*_ in the analytical approach by *H*_*c,tot*_ = 2*H*_*c*_, see Equation 1b. Due to this convention, the coat spontaneous curvature parameter *C*_*c*_ used in our continuum model must be scaled accordingly (by a factor of two) when making quantitative comparisons with geometrical models that define the mean curvature as the arithmetic average *H*_*c,tot*_*/*2.

The spontaneous curvature *C*(*ϕ*_*S*_), bending rigidity *κ*(*ϕ*_*S*_), and effective surface tension *σ*_*S*_(*ϕ*) interpolate as:

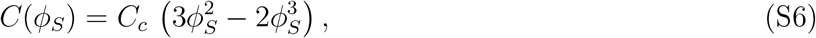

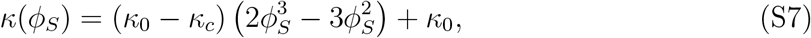

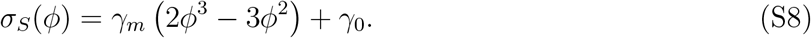

The term *σ*_*S*_(*ϕ*) specifically accounts for the wetting forces by differentiating between membrane-droplet adhesion and membrane-cytosol interfacial tension *γ*_*m*_ and *γ*_0_, respectively. We set *γ*_0_ = 3.2 pN/nm. Because the membrane is locally incompressible, a uniform shift of *σ*_*S*_ changes *E*_*S*_ only by a constant and leaves all forces unchanged; *γ*_0_ is therefore set not by measurement but by the requirement that *σ*_*S*_(*ϕ*) ≥ 0 everywhere. The geometrical model does not impose local incompressibility and so uses the physical membrane tension *σ* directly.

#### Force balance and incompressibility

The coupling between the fluid and the membrane is realized via a stress balance condition. The jump [**S** · **n**]_*S*_ in fluid stress **S** across the interface is balanced by the intrinsic membrane forces **f**_*S*_, where **n** denotes the normal vector of *S* pointing towards the external fluid. Crucially, lipid membranes exhibit local area incompressibility. We enforce the constraint ∇_*S*_ · **u** = 0, taking into account the surface divergence ∇_*S*_ ·, by introducing a local Lagrange multiplier *ξ* that provides the necessary additional surface tension. The total force balance at the membrane reads:

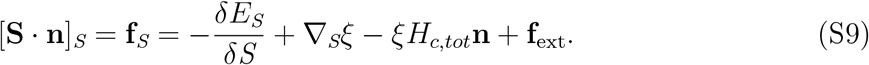

#### External forces

To capture the specific physical environment of yeast endocytosis, the term **f**_ext_ includes two contributions:

- **Turgor pressure:** A constant normal force density pointing outwards, **f**_turgor_ = *P*_turgor_ **n**.
- **Cell wall interaction:** A repulsive penalty force preventing non-physical outward bulging, modeled as:

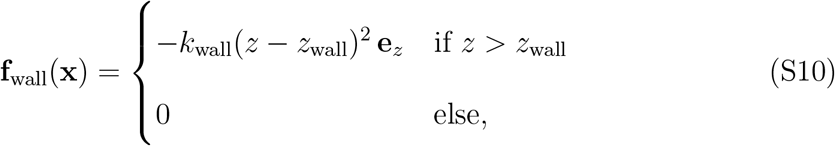

where the wall is located perpendicular to the *z*-coordinate at *z* = *z*_wall_.

### Evolution of the protein coat

The dynamics of the protein coat on the evolving membrane are modeled using a conservative surface Cahn-Hilliard equation for the order parameter *ϕ*_*S*_. Since we strictly enforce local area incompressibility of the lipid bilayer (∇_*S*_ · **u** = 0), the surface dilation term vanishes. The advective transport of the coat is governed by

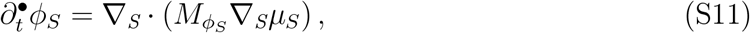

where 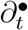 denotes the material time derivative following the membrane flow, and *µ*_*S*_ is the surface chemical potential. We note that surface diffusion merely serves to stabilize advection and keep a fixed interface profile. To satisfy the condition of a constant coated area and prevent artificial deviations of the coated region due to numerical effects during large geometric deformations, we add an explicit area-conservation term at the discretization level (see numerical details below).

### Numerical implementation

The system of coupled partial differential equations is solved using the finite element toolbox AMDiS,^66,67^ which is based on the DUNE framework.^68^ The domain deformation is handled using an Arbitrary Lagrangian-Eulerian (ALE) approach. Time discretization is performed using a semi-implicit scheme to ensure stability. We utilize an axisymmetric coordinate system to reduce computational cost while retaining the full 3D geometry of the invagination. The membrane is modeled as locally incompressible via the aforementioned Lagrange multiplier method. At the outer boundary of the membrane patch, we apply a free-slip condition (relaxing the zero normal velocity constraint) to allow lipid material to flow into the computational domain, mimicking the recruitment of membrane area from the surrounding plasma membrane reservoir.

### Numerical discretization

The numerical framework used here extends the model introduced by Mokbel et al.,^54^ which combines a diffuse-interface description of the bulk fluids with a sharp-interface ALE method for the deformable membrane.

#### Time discretization and ALE framework

We consider the time interval [0, *T*] partitioned into uniform time steps of size *τ*, with discrete time points *t*_*n*_ = *nτ*. To accommodate the large topological deformations of the membrane *S*(*t*) (such as endocytic invagination) while preserving the quality of the bulk mesh, we utilize the ALE method. The computational domain Ω(*t*) is partitioned into an intracellular inner domain Ω_in_(*t*) and an extracellular outer domain Ω_out_(*t*), discretized by a moving mesh that conforms explicitly to the membrane interface. Because the phase-separated droplet forms inside the cell, the bulk phase field *ϕ* is strictly evaluated within Ω_in_(*t*).

The mesh velocity **w** is constructed to track the membrane interface. To prevent numerical grid degeneracies during large deformations, the grid movement of the membrane is restricted to the normal component of the fluid velocity, i.e., **w**|_*S*_ = (**u**|_*S*_ · **n**)**n**. In the bulk subdomains, **w** is computed as the harmonic extension of the boundary velocity, governed by Δ**w** = 0, to prevent mesh entanglement.

To formulate a stable and efficient scheme, we employ a semi-implicit (IMEX) Euler time discretization. The time derivative of any spatial field *f* is treated via the ALE material derivative. In the discrete setting, this introduces an explicit convective transport term relative to the mesh movement, (**u** − **w**) · ∇*f*, in the bulk equations. The geometry is updated explicitly after each time step via Ω^*n*+1^ = *{***x** + *τ* **w**^*n*^(**x**) | **x** ∈ Ω^*n*^*}*.

### Finite element spaces and pressure discontinuity

Let *T*_*h*_ be a conforming simplicial triangulation of the bulk domain Ω, strictly partitioned into the inner domain *T*_*h*,in_ and the outer domain *T*_*h*,out_. The restriction of this triangulation to the membrane surface *S* forms a (*d* − 1)-dimensional mesh *S*_*h*_ = *T*_*h*,in_ ∩ *T*_*h*,out_.

A critical requirement of modeling membranes subject to surface tension and Helfrich bending forces is the exact resolution of the pressure jump across the interface, dictated by the generalized Young-Laplace law. Standard continuous finite elements for pressure yield sub-optimal 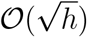 convergence and spurious parasitic currents. Therefore, we define split pressure spaces, effectively extending the Taylor-Hood finite element space with additional degrees of freedom at the interface:

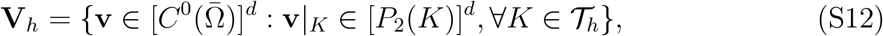

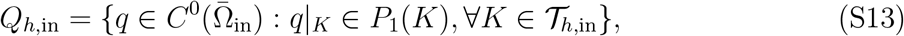

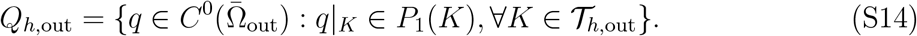

The bulk phase field *ϕ* and chemical potential *µ* are discretized using standard second-order continuous elements 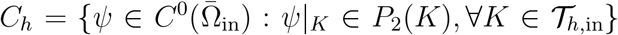, reflecting their restriction to the intracellular space. Geometric variables on the membrane, including the spatial coordinate vector 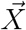, the generalized curvature vector 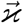 (see below), the membrane force **f**_*S*_, the surface phase field *ϕ*_*S*_, and the Lagrange multiplier *ξ*, are discretized using first-order Lagrange elements on the surface:

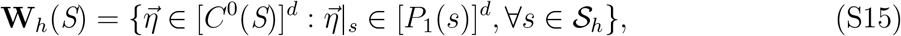

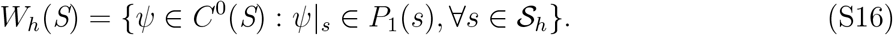

### Fully discrete weak formulation

The Helfrich bending energy introduces fourth-order geometric derivatives, which cannot be discretized directly using *C*^0^-continuous finite elements. To bypass this restriction, we utilize a mixed finite element formulation initially introduced by Dziuk^69^ and expanded for multi-phase membranes by Barrett, Garcke, and Nürnberg.^70^ We introduce a generalized curvature vector 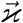 (acting as the bending moment vector) and the total membrane force **f**_*S*_ as independent primary variables.

The coupled system is solved monolithically in each time step. We search for the fluid velocity **u**^*n*+1^ ∈ **V**_*h*_, pressures 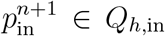 and 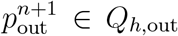, the bulk phase field *ϕ*^*n*+1^, *µ*^*n*+1^ ∈ *C*_*h*_, the generalized curvature vector 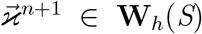, the membrane force 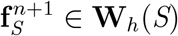, the surface tension Lagrange multiplier *ξ*^*n*+1^ ∈ *W*_*h*_(*S*), and the surface order parameter 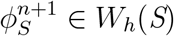

#### Hydrodynamics and bulk phase field

The incompressible Navier-Stokes equations are coupled to the Cahn-Hilliard equations via a semi-implicit capillary stress tensor acting within the intracellular domain. Note that while we formulate the equations for constant density *ρ*, we consider the system in the Stokes flow regime in this study; hence, the non-linear convective Navier terms are omitted in the numerical solutions. Find 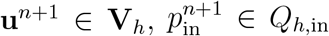, and 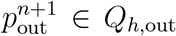 such that for all test functions **v** ∈ **V**_*h*_, *q*_in_ ∈ *Q*_*h*,in_, and *q*_out_ ∈ *Q*_h,out_:

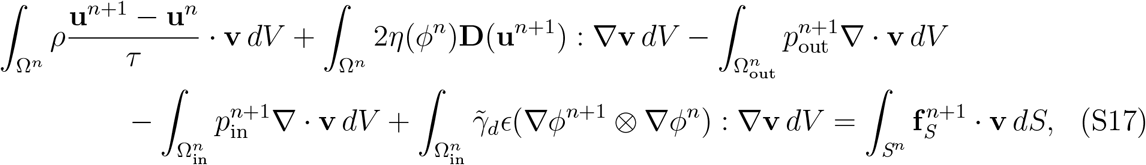

subject to the divergence-free constraints 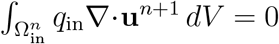 and 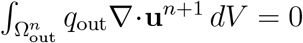. Here, 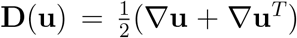 is the rate-of-deformation tensor, and the capillary stress is linearized semi-implicitly.

Simultaneously, find *ϕ*^*n*+1^, *µ*^*n*+1^ ∈ *C*_*h*_ such that for all test functions *ψ*_1_, *ψ*_2_ ∈ *C*_*h*_:

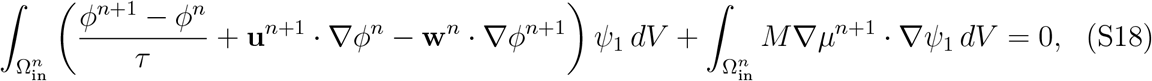

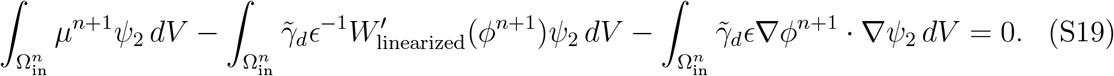

The double-well potential derivative is treated via the Taylor expansion 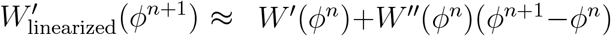 to guarantee stability, and the phase-field Laplacian is integrated by parts.

#### Geometric evolution and variable bending rigidity

To handle the spatially varying material properties induced by the protein coat, the variable 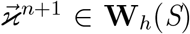 implicitly incorporates the local bending rigidity 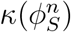 and the spontaneous curvature 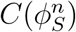. Find 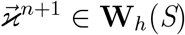 such that for all 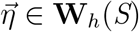:

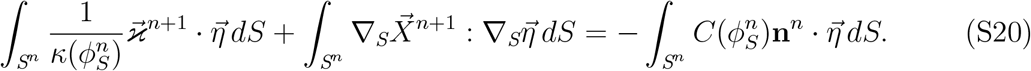

The spatial coordinate vector 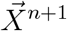 is updated implicitly via the relation 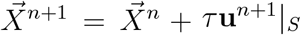. The term 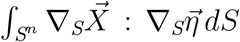 represents the weak form of the surface Laplacian 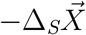, which corresponds to the geometric curvature vector. Consequently, Eq. (S20) effectively enforces 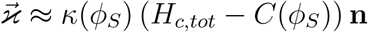.

#### Membrane force balance and incompressibility

The total force exerted by the membrane on the fluid, 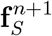, is assembled from the variation of the Helfrich energy, tension variations, and external cellular forces. Following the multi-phase membrane methodology from Barrett, Garcke, and Nürnberg,^70^ the implicit weak formulation precisely accounts for metric variations. Find 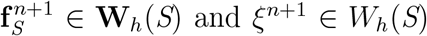 and *ξ*^*n*+1^ ∈ *W*_*h*_(*S*) such that for all 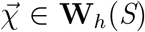 and

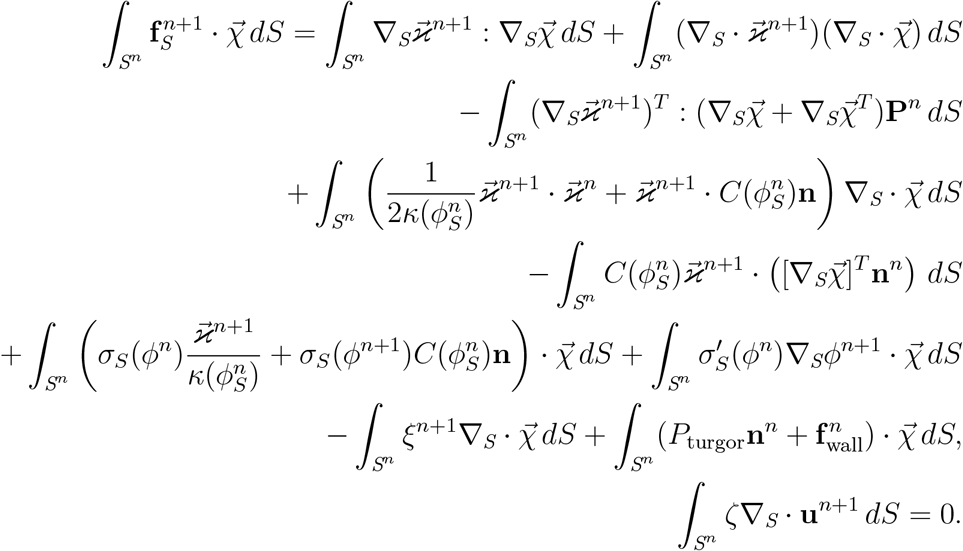

Here, **P** = **I** − **n** ⊗ **n** denotes the surface projection operator. The last equation strictly enforces local area incompressibility (∇_*S*_ · **u** = 0).

#### Surface phase field (protein coat)

To discretize the transport equation on the evolving membrane, we utilize the ALE framework. The time derivative along the moving mesh introduces a geometric source term proportional to the surface divergence of the grid velocity **w**. Because our mesh moves purely in the normal direction, i.e., **w** = (**u** · **n**)**n**, this surface dilation is strictly determined by the local curvature *H*_*c,tot*_, reading ∇_*S*_ · **w** = *H*_*c,tot*_(**u** · **n**). In our fully discrete scheme, we express this geometric term using the generalized curvature vector 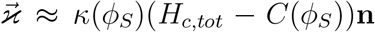. Rearranging this relation yields 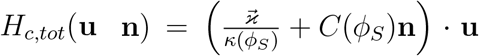, which provides the exact substitution used in the advection integral. This ensures strict numerical consistency between the geometric transport and the mechanical bending forces.

Additionally, to strictly enforce the constant coat area constraint and counteract numerical dissipation, an explicit area-correction term proportional to |∇_*S*_*ϕ*_*S*_| is added. Instead of a constant parameter, the prefactor 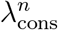 is evaluated dynamically in each time step to penalize relative deviations from the initial coat area *A*_0_. The fully discrete scheme reads: Find 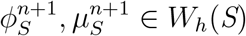 such that for all test functions *ζ, ψ* ∈ *W*_*h*_(*S*):

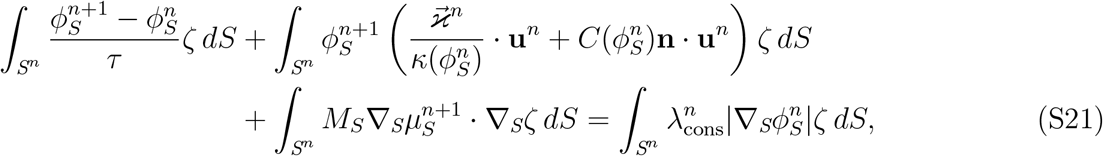

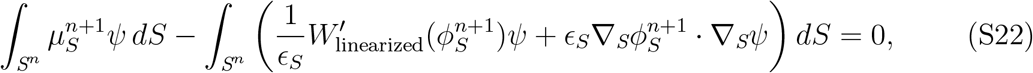

where the dynamic penalty factor is defined as 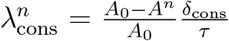. Here, 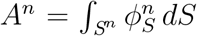 is the total coated area at the current time step, *A*_0_ is the target area, *δ*_cons_ is a user-defined scaling parameter, and *τ* is the time step size. Moreover, note that the chemical potential is chosen here in an analogous manner to Equation S19.

The resulting monolithically coupled linear system is assembled and solved using a direct sparse solver (e.g., UMFPACK) provided by the Dune-ISTL library within the AMDiS ecosystem.

### Finite element model parameters

In addition to the values in Table 1, we choose the following parameters in the numerical solutions: *M* = 5.328 · 10^−20^ m^3^s/kg, *ϵ* = 9 · 10^−10^ m, *M*_*S*_ = 3.33 · 10^−15^ m^3^/s, *ϵ*_*S*_ = 1 · 10^−9^ m, Δ*t* = 4 · 10^−9^ s, and *η* = 0.05 Pa·s (constant viscosity). The mesh size is ≈ 1.36 · 10^−9^ m in the membrane-and droplet interface regions and ≈ 2.23 · 10^−8^ m in the bulk regions. The cell wall parameter *k*_wall_ can theoretically be set to any value and has been calibrated so that it is large enough to ensure sufficient repulsion while not causing numerical instabilities.

**Table 1.** Parameters chosen in models. All analogous parameters among both the finite-element model and geometrical model were set to be identical except for κ_c_, which was tuned to recapitulate equivalent membrane dynamics between models (Figure S2). γ_m_ was not measured independently; it was set to 0.32γ_d_, which fixes the droplet contact angle at θ_d_ ≈ 71^°^ through Young’s equation (Equation S29), and is therefore rescaled with any change to γ_d_. C_c_ was set so that the preferred radius of curvature of the coat matches the final vesicle radius.

| Description | Symbol | Value | Units | References |
| --- | --- | --- | --- | --- |
| Membrane tension | $\sigma$ | 0.2 | pN/nm | 40–43 |
| Droplet-cytosol surface tension | $\gamma_d$ | 10 | pN/nm | 44,45 |
| Droplet-membrane adhesion energy density | $\gamma_m$ | 3.2 | pN/nm | Model choice (see caption) |
| Coat bending rigidity (finite-element model, geometrical model) | $\kappa_c$ | (3173, 1500) | pN · nm | 40,46–48 |
| Coat spontaneous curvature | $C_c$ | 0.05 | nm<sup>−1</sup> | Model choice (see caption) |
| Coat surface area | $S_c$ | $4\pi 20^2$ ($\approx 5 \cdot 10^3$) | nm<sup>2</sup> | 48–50 |
| Bare membrane bending rigidity | $\kappa_o$ | 160 | pN · nm | 40,48 |
| Bare membrane spontaneous curvature | $C_o$ | 0 | nm<sup>−1</sup> | 40 |
| Turgor pressure | $P$ | 0.5 | pN/nm<sup>2</sup> | 50–53 |

Similarly, *δ*_cons_ has been calibrated to be large enough to maintain the coated area without causing numerical instabilities.

### Construction of geometrical model

We complement the finite-element model with a simplified geometrical model in order to analyze the generalizability of the numerical solutions.

#### System geometry and constraints

The model consists of a spherical cap droplet wetting a spherical cap membrane invagination from a flat disk membrane base (Figure S1). All relevant geometrical quantities are defined either as constants or as functions of the relative invagination distance *h*. The droplet has a constant volume *V*_*d*_, constant contact angle *θ*_*d*_, *h*-dependent cytosolic contact area *S*_*d*_, and *h*-dependent membrane contact area *S*_*m*_. The membrane has a constant clathrin-coated surface area *S*_*c*_, *h*-dependent displaced cytoplasmic volume *V*_*c*_, and *h*-dependent cytosolic contact area *S*_*o*_.

Using the definition of a spherical cap, *h* is formally defined as

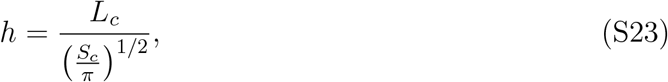

where *L*_*c*_ is the height of the invagination spherical cap in units of nm.

The spherical cap defines *S*_*d*_ as:

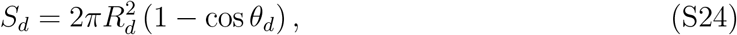

where *R*_*d*_ is the radius of the spherical cap consisting of the combined droplet and membrane invagination:

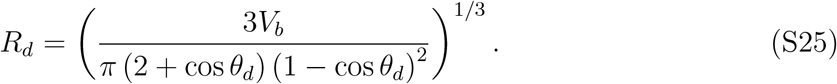

*V*_*b*_ is the combined droplet and membrane invagination volume:

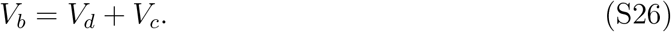

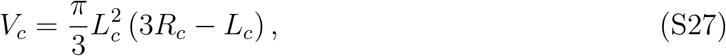

where *R*_*c*_ is the radius of the coated membrane invagination spherical cap:

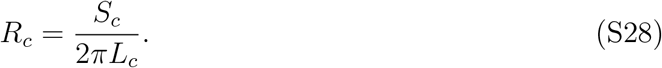

With the definition for *θ*_*d*_ provided by Young’s equation

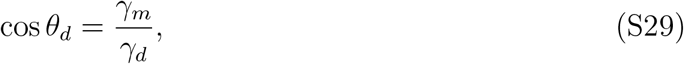

we can substitute and simplify to express *S*_*d*_ in terms of *h*:

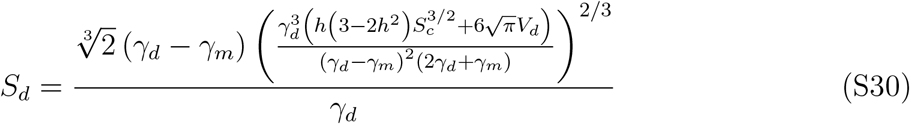

For easier comparison to other geometry equations, we rearrange this to pull out a pre-factor:

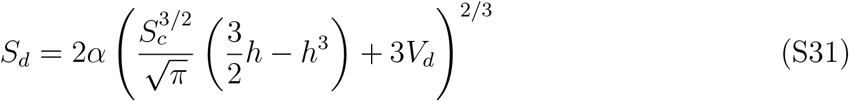

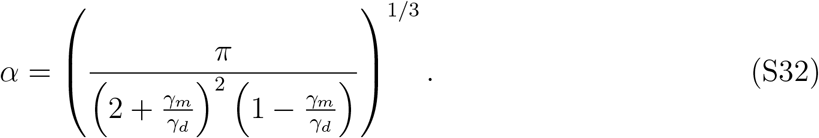

We take the derivative with respect to *h* in order to analyze the rate of expansion or contraction of this area at a given point in the pathway:

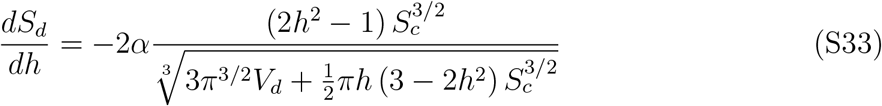

Next, the droplet-membrane contact area *S*_*m*_ is determined by:

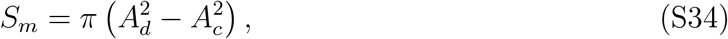

where *A*_*c*_ is the area of the plane of the base of the membrane invagination and *A*_*d*_ is the combined droplet and membrane base plane area. These are given by

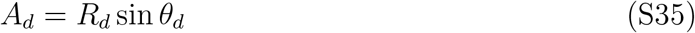

and

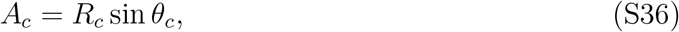

where *θ*_*c*_ is the contact angle of the membrane invagination spherical cap with the base membrane:

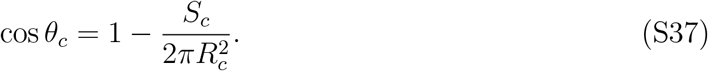

Once again, we substitute and refactor to express *S*_*m*_ in terms of *h*:

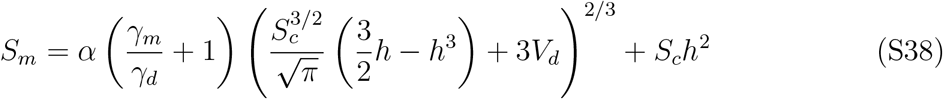

and solve the derivative:

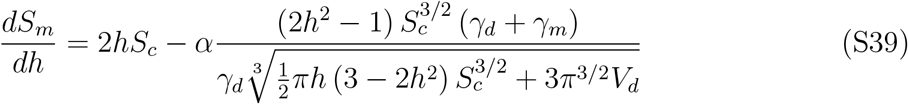

As we are interested in comparing the rates of expansion for each interfacial area at the start of the pathway, we evaluate 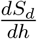 (Equation S33) and 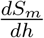 (Equation S39) at *h* = 0:

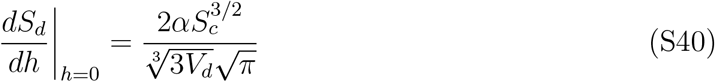

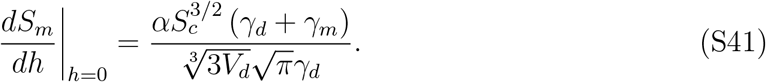

We substitute these two equations to result in Equation 3.

### Free energy and forces

As introduced in Equation 1, the free energy *F* of the geometrical model is

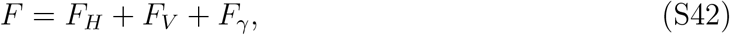

which breaks down to

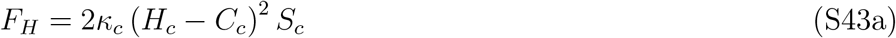

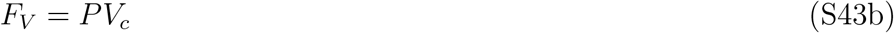

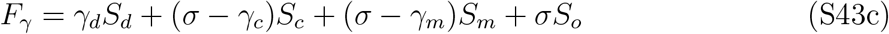

The bending energy *F*_*H*_ depends on the bending rigidity of the coated membrane *κ*_*c*_, its mean curvature *H*_*c*_, its protein-induced spontaneous curvature *C*_*c*_, and its area *S*_*c*_. The pressure energy *F*_*V*_ depends on the osmotic pressure across the membrane *P* and the volume displaced by the coated invagination *V*_*c*_. The surface energy *F*_*γ*_ depends on the membrane tension *σ* and the sum of all membrane segments *S*_*c*_, *S*_*m*_, and *S*_*o*_, the droplet-cytosol interfacial area *S*_*d*_ and associated tension *γ*_*d*_, droplet-coated membrane interfacial area *S*_*c*_ and associated tension *σ* − *γ*_*c*_, droplet-bare membrane interfacial area *S*_*m*_ and associated tension *σ* − *γ*_*m*_, and cytosol-bare membrane interfacial area *S*_*o*_ and associated tension *σ*.

Previously published models of yeast endocytosis write the bending energy with the total curvature *c*_1_ + *c*_2_ and a prefactor 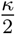.^48,50^ That form is identical to Equation S43a under *c*_1_ + *c*_2_ = 2*H*, so their *κ* equals *κ*_*c*_ and their spontaneous curvature equals 2*C*_*c*_. All literature values quoted in this work are converted to our mean-curvature convention.

As we constrain a constant coat surface area (Δ*S*_*c*_ = 0), the change in free energy over the pathway Δ*F* = *F* − *F* |_*h*=0_ simplifies to:

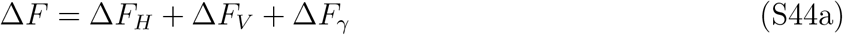

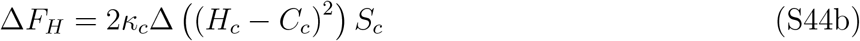

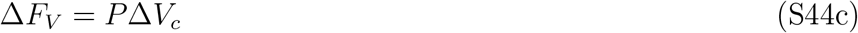

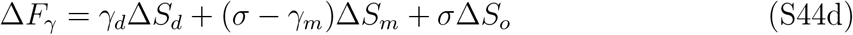

We are interested in the droplet-mediated energy contribution Δ*F*_*d*_ by isolating the droplet terms:

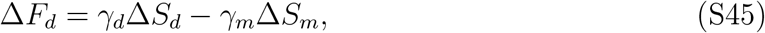

composed of the free energy contribution from the droplet-cytosol interface *γ*_*d*_Δ*S*_*d*_ and the free energy contribution from the droplet-membrane interface *γ*_*m*_Δ*S*_*m*_. Substituting into Equation S31 and Equation S38:

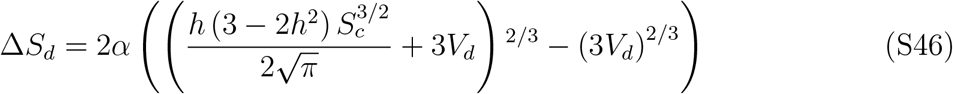

and

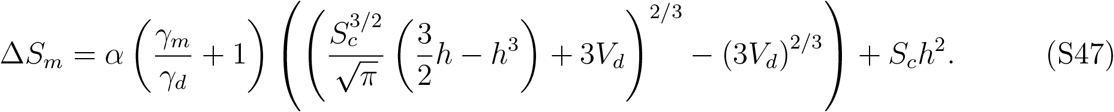

We calculate the force from the droplet by taking the derivative of *F*_*d*_ with respect to *h*:

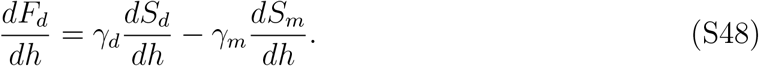

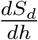 and 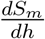 are defined in Equation S33 and Equation S39, respectively.

Forces reported in units of pN are derivatives with respect to the physical invagination distance *L*_*c*_, not the dimensionless *h*. Equation S23 relates the two by a constant factor:

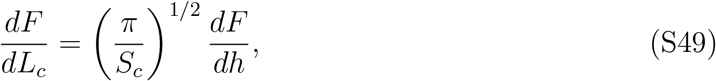

so that the droplet capillary force is

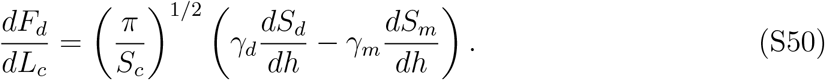

Conversely, the droplet-independent change in energy Δ*F*_*e*_ is:

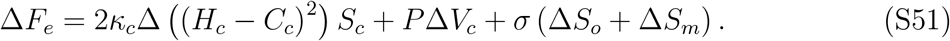

The mean curvature of the coated membrane invagination is given by 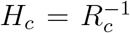, with *R*_*c*_ defined in Equation S28.

The derivative provides the droplet-independent force:

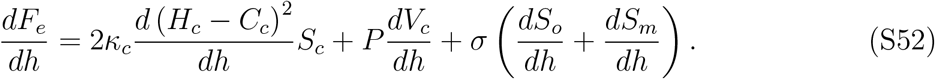

Substituting Equation S28 into 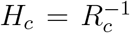 gives 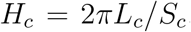, so the coat curvature-generating force is

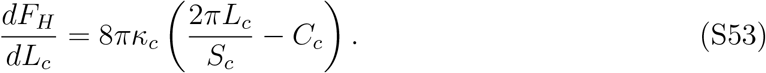

The turgor pressure term in Equation S52 acts on an effective area constrained by the fixed coated membrane area, which determines how the displaced volume *V*_*c*_ changes with invagination depth. Combining Equation S27 with Equation S28 sets the turgor pressure force as

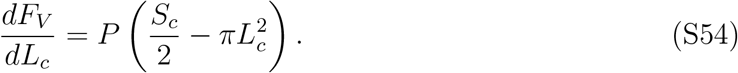

The membrane tension term in Equation S52 needs no separate expression for *S*_*o*_. Both base regions stay flat and the coat area is fixed, so *S*_*o*_ + *S*_*m*_ is the flat annulus outside the base of the coated cap, and only its inner radius moves. Combining Equation S36 and Equation S37 with Equation S28 gives

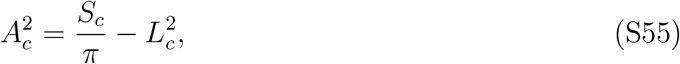

so the flat base area grows at a rate set by *L*_*c*_ alone, with no droplet dependence:

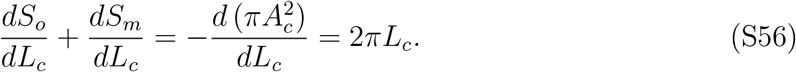

The membrane tension force is therefore

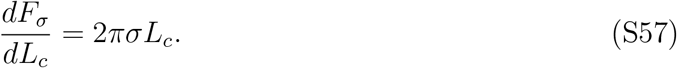

### Kinetic stability analysis

In order to analytically determine the boundary between stalling and invaginating initial conditions, we calculate the parameter values necessary to satisfy 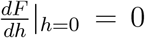. This construction allows us to plot two-dimensional state diagrams by solving for one of the axis parameters.

The initial force in the system 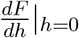 in the fully expanded form in terms of *h* is:

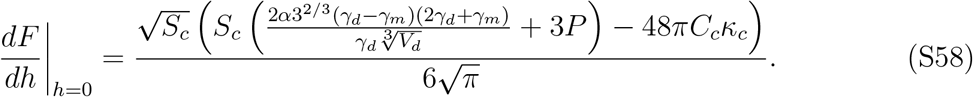

For Figure 4 A and D and Figure S2 A, we isolate *V*_*d*_ in the 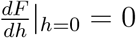 condition in order to solve the critical volume 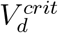 as other parameters vary:

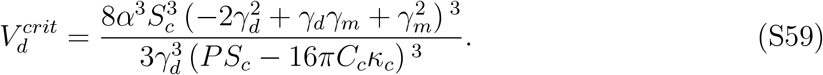

Figure 4 D plots the two sides of this balance separately at the flat initiation state, in units of pN. The droplet capillary force is Equation S50 evaluated with Equation S40 and Equation S41. The non-droplet force is the sum of Equation S53 and Equation S54 at *L*_*c*_ = 0, where the membrane tension contribution (Equation S57) vanishes. It is plotted with the opposite sign, so that the two curves cross at 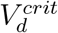 rather than sum to zero:

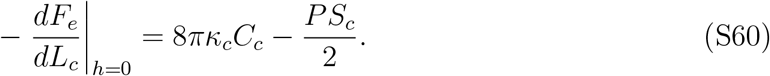

For the reference parameters this evaluates to 6.3 · 10^2^ pN.

**Figure S1:**
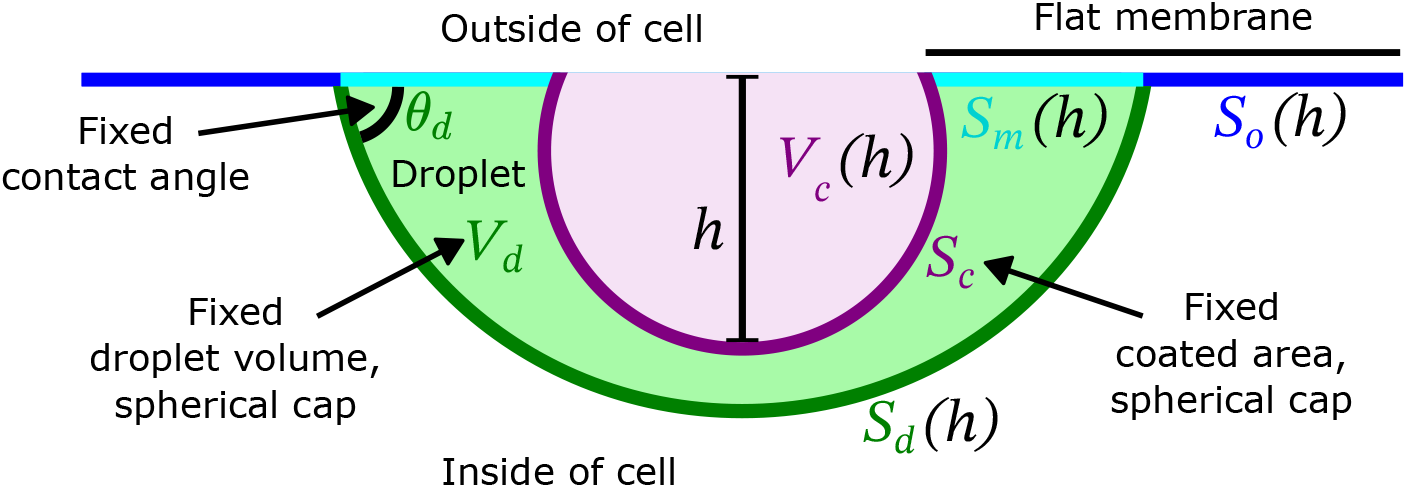
Cartoon schematic of geometrical model. A protein droplet with constant volume *V*_*d*_ partially wets a patch of the plasma membrane with a contact angle *θ*_*d*_ set by the balance of surface tensions. It has a contact area with surrounding cytosol *S*_*d*_. The plasma membrane at the site of endocytosis is divided into three regions: the surface contacting the cytosol *S*_*o*_, the surface contacting the droplet *S*_*m*_, and the clathrin-coated surface *S*_*c*_. *S*_*o*_ and *S*_*m*_ remain flat surfaces. *S*_*c*_ can transition from the flat initiation stage to the final spherical scission stage through a spherical cap intermediate with constant surface area. With the aforementioned geometrical constraints, the entire system is defined by a single reaction coordinate *h*, the normalized membrane invagination height. All surface areas and volumes that depend on *h* are labeled as functions of *h* in the schematic cartoon: *S*_*o*_(*h*), *S*_*m*_(*h*), *S*_*d*_(*h*), and *V*_*c*_(*h*).

**Figure S2:**
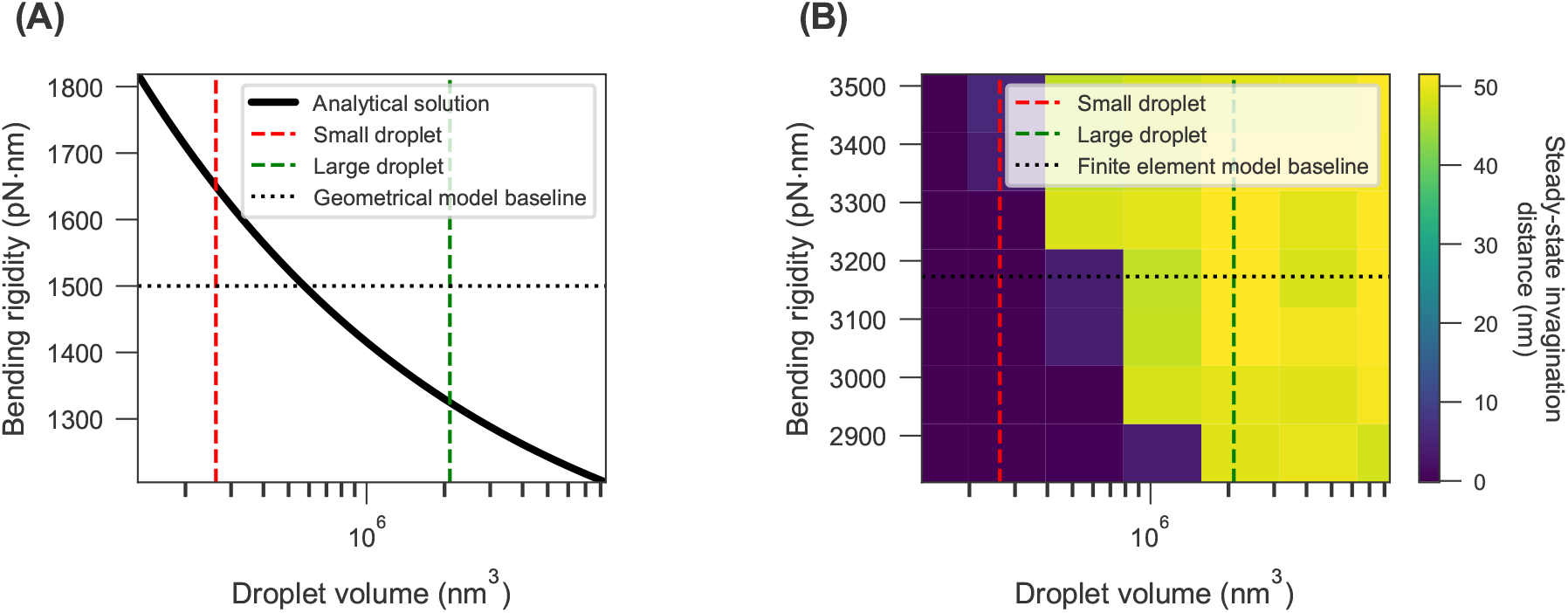
Side-by-side theory vs. numerical solution calibration comparison. (A) Analytical solution of contour line where 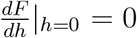 (Equation 4) over varying droplet volume and coat bending rigidity. At the model’s fixed value of *κ*_*c*_ = 1500 pN nm, the system can transition from stalled to unstalled within a realistic droplet growth regime. (B) With the same parameters as in (A), the finite-element model was evaluated over an array of bending rigidities in order to calibrate a value *κ*_*c*_ = 3173 pN nm that recapitulates the droplet volume-dependent stalling behavior from (A). In both panels, red and green dashed lines mark the small and large droplet volumes of Figure 4 B (*V*_*d*_ ≈ 2.6 · 10^5^ and ≈ 2.1 · 10^6^ nm^3^).

**Figure S3:**
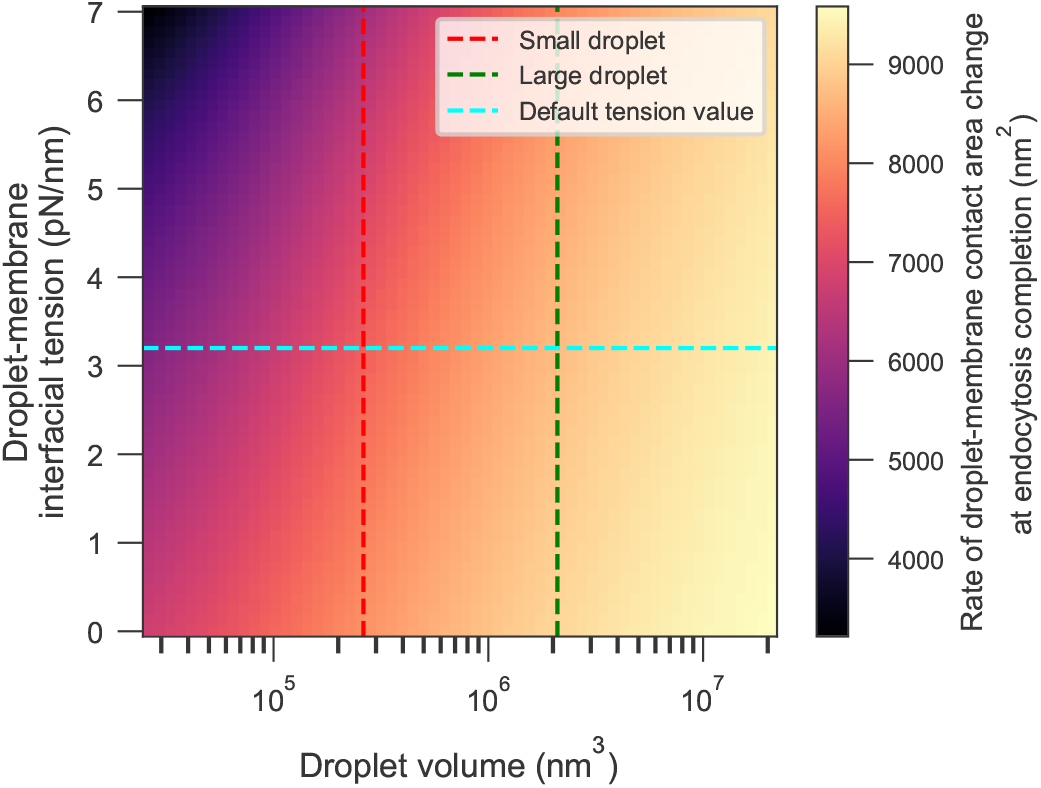
Late endocytosis droplet-membrane contact area rate of change over relevant parameter range. The color scale represents 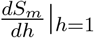, which is positive for the entire parameter range that is relevant to endocytosis. This indicates that a droplet is always generating an assistive force at the end of the invagination trajectory in the geometrical model. The red and green dashed lines mark the small and large droplet volumes of Figure 4 B (*V*_*d*_ ≈ 2.6 · 10^5^ and ≈ 2.1 · 10^6^ nm^3^), and the cyan dashed line the reference droplet-membrane interfacial tension *γ*_*m*_ = 3.2 pN/nm.

**Figure S4:**
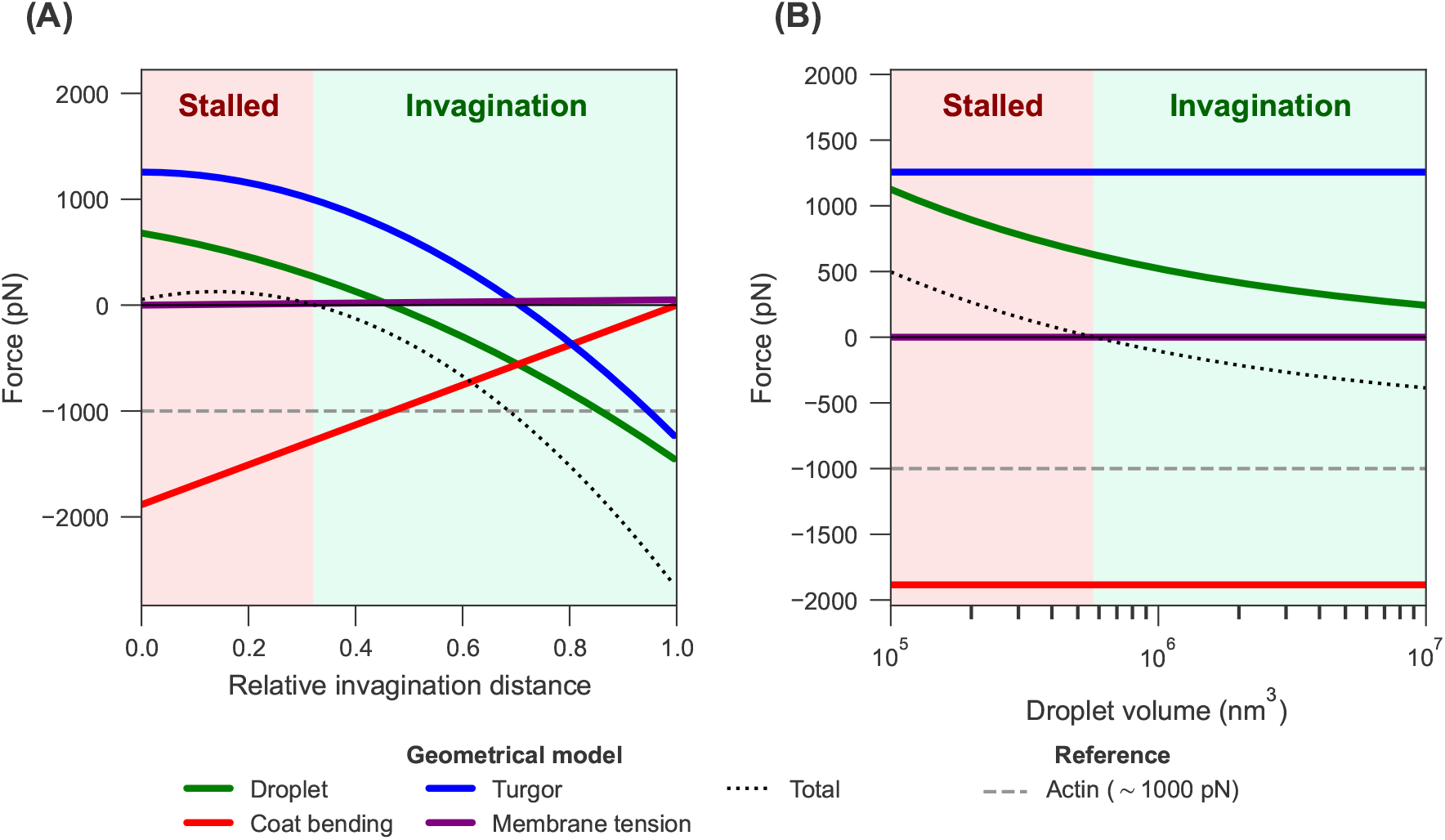
Comparison of individual force contributions in the geometrical model. Forces are computed as the derivative of the free energy in Equation 1 with respect to the physical invagination distance *L*_*c*_ (Equation S23), where *h* = 1 corresponds to 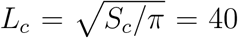 at the reference coat surface area. (A) Individual force components vs. relative invagination distance *h* across the full internalization pathway. Plotted are the droplet capillary force, the coat curvature-generating force, the turgor pressure force, the membrane tension force, and total force, which is the sum of the model terms. Sign convention: positive forces oppose invagination; negative forces drive invagination. The horizontal dashed reference line indicates the force from actin network assembly transmitted to the plasma membrane at a budding yeast CME site, estimated at approximately 1000 pN.^59,60^ Note that actin is not a component in the model and it is not included in the line for total force. (B) The same force components at the flat initiation state (*h* = 0) as a function of droplet volume *V*_*d*_. The non-droplet components are independent of *V*_*d*_; the droplet force decreases with *V*_*d*_, and the total crosses zero at the critical droplet volume. Reference parameters as in Table 1 unless otherwise indicated.

**Movie S1. Membrane invagination without a droplet**. Three-dimensional rendering of the finite element solution shown in Figure 3 A (top). The coated membrane patch (purple) bends away from the flat membrane base (teal) and forms a nascent vesicle. No droplet is present (*V*_*d*_ = 0).

**Movie S2. A droplet stalls membrane bending**. Three-dimensional rendering of the finite element solution shown in Figure 3 A (bottom), for a droplet of volume *V*_*d*_ ≈ 2.6 · 10^5^ nm^3^ (green). The coated patch (purple) reaches a shallow stationary shape and does not invaginate further.

**Movie S3. Droplet volume sets the energy barrier to invagination**. Geometrical model solutions as the droplet volume *V*_*d*_ increases from 2.6 · 10^5^ to 2.1 · 10^6^ nm^3^. (A) Cross-section at the local free energy minimum, showing bare membrane (blue), coat (magenta), droplet-membrane interface (cyan), and droplet-cytosol interface (green). (B) Change in droplet-cytosol interfacial energy (green line), droplet-membrane interfacial energy (cyan line), and total droplet energy (black line) over relative invagination distance *h*. (C) Change in total free energy Δ*F* over *h*. The red point marks the local minimum drawn in (A). The barrier vanishes as *V*_*d*_ passes the critical volume of Figure 4 D.

**Movie S4. Droplet growth releases the stall**. Three-dimensional rendering of the finite element solution shown in Figure 4 E. The droplet (green) grows at a constant rate. The coated membrane (purple) stays stalled while the droplet is small, then rapidly invaginates once the droplet passes the critical volume.

